# High Spatial-Resolution Imaging of the Dynamics of Cuticular Lipid Deposition During Arabidopsis Flower Development

**DOI:** 10.1101/2020.11.17.387241

**Authors:** Liza Esther Alexander, Jena S. Gilbertson, Bo Xie, Zhihong Song, Basil J. Nikolau

**Affiliations:** Roy J. Carver Department of Biochemistry, Biophysics and Molecular Biology, Iowa State University, Ames, Iowa, 50011; Center for Metabolic Biology, Iowa State University, Ames, Iowa 50011

**Author notes:** **Corresponding Author:** Basil J. Nikolau. Roy J. Carver Department of Biochemistry, Biophysics and Molecular Biology, Iowa State University, Ames, IA, 50011, USA. Jena S. Gilbertson, Illinois College of Optometry, 3241 S Michigan Ave, Chicago, IL-60616. Bo Xie, Office of Intellectual Property and Technology Transfer, Economic Development Core Facility, 1805 Collaboration Place, Suite 2100, Iowa State University, Ames, IA-50010. Email address Zhihong Song, Office of Pharmaceutical Quality, Center for Drug Evaluation and Research, U.S. Food and Drug Administration, 10903 New Hampshire Avenue, Silver Spring, MD-20993.

## Abstract

The extensive collection of *glossy* (*gl*) and *eceriferum* (*cer*) mutants of maize and Arabidopsis have proven invaluable in dissecting the branched metabolic pathways that support cuticular lipid deposition. This branched pathway integrates the fatty acid elongation-decarbonylative branch and the fatty acid elongation-reductive branch that has the capacity to generate hundreds of cuticular lipid metabolites. In this study a combined transgenic and biochemical strategy was implemented to explore and compare the physiological function of three homologous genes, *Gl2, Gl2-like* and *CER2* in the context of this branched pathway. These biochemical characterizations integrated new extraction-chromatographic procedures with high-spatial resolution mass spectrometric imaging methods to profile the cuticular lipids on developing floral tissues transgenically expressing these transgenes in wild-type or *cer2* mutant lines of Arabidopsis. Collectively, these datasets establish that both the maize *Gl2* and *Gl2-like* genes are functional homologs of the Arabidopsis *CER2* gene. In addition, the dynamic distribution of cuticular lipid deposition follows distinct floral organ localization patterns indicating that the fatty acid elongation-decarbonylative branch of the pathway is differentially localized from the fatty acid elongation-reductive branch of the pathway.

## INTRODUCTION

The multicellular nature of plants has provided challenges to deciphering the genetic and biochemical mechanisms that regulate extracellular cuticular lipid biogenesis. These lipids are produced by the epidermal cell layer that accounts for only about 10% of the cellular population of the aerial organs of plants (Jellings and Leech, 1982). Additionally, this biological process is physically distributed among different compartments at the cellular and subcellular levels (cytoplasm, endoplasmic reticulum, and plasma membrane of epidermal cells). The barriers associated with this complexity have been partially overcome with forward genetic approaches that use *eceriferum* (*cer*) mutants of Arabidopsis (Koornneef et al., 1989), *glossy* (*gl*) mutants of maize (Schnable et al., 1994) and tomato (Leide et al., 2007; Vogg et al., 2004) and *wax crystal-sparse leaf* (*wsl*) mutants of rice (Wang et al., 2017; Yu et al., 2008), which has led to the molecular identification and characterization of genes involved in cuticle deposition. Many examples indicate however, that the isolation of the causative gene that gives rise to the defect in cuticular lipid deposition is only the starting point for detailed understanding of the functionality of the isolated gene in the context of cuticle biosynthesis (Bernard and Joubès, 2013; Post-Beittenmiller, 1996; Samuels et al., 2008).

These complexities can be partially overcome by applying detailed proteomic and/or metabolomics characterizations, comparing mutant and wild-type plants. However, a particular limitation with these experiments is the fact that these techniques are generally performed on tissue extracts that homogenize tissues and cells that are at different developmental stages, and thereby losing *in situ* localization and developmental states of the cell population. Technologies that make use of “reporter” interactions can visualize gene expression at high spatial scale resolution; for example *in situ* nucleic acid hybridization and antibody reactivity (Griffin et al., 1998; Küpper et al., 2007; McFadden, 1995), or genetic fusion with fluorescent reporters (Chalfie et al., 1994; Gallagher, 1992; Koo et al., 2007; Ow et al., 1986). Thus, these technologies produce data concerning the spatial distribution of macromolecules and generate data that provide indirect insights of the *in vivo* flow of intermediates of metabolism through metabolic networks in and among different cells. Moreover, data generated by these techniques are not necessarily dynamic. This gap in the biochemical characterization pertaining to the nature and regulation of metabolism can be partially overcome with mass spectrometric imaging (MSI) (McDonnell and Heeren, 2007).

MSI is a rapidly growing technology known for its ability to provide high spatial resolution data on the cellular location of metabolites, detecting extremely small amounts of metabolites (i.e., high sensitivity), and providing chemical identification data concerning the detected analyte. Additionally, among the several molecular ionization techniques that are used in MSI, the use of diverse ionization technologies as in the case of matrix assisted laser desorption ionization (MALDI)-MSI, further enhances the abilities of this technology. Hence, it is now possible to analyze the spatial distribution of a variety of different classes of compounds (e.g., lipids, proteins, and small molecules) in both plant and animal tissues (Amstalden van Hove et al., 2011; Angel and Caprioli, 2013; Jungmann and Heeren, 2012; Lee et al., 2012; Svatoš, 2010).

As exemplary application of this imaging technology, the focus of this study was the characterization of two homologous maize genes, *Gl2* and *Gl2-like*, whose specific biochemical functions are still unclear, but both genes are known to impact cuticular lipid deposition (Bianchi, 1975;Tacke et al., 1995; Alexander et al., 2020). *Gl2* was initially identified as a product of a forward genetic screen (Hayes and Brewbaker, 1928) and subsequently, following its molecular characterization (Tacke et al., 1995) it led to the identification of the homologous *Gl2-like* gene (Alexander et al., 2020). Similar to *Gl2*, the *CER2* gene was initially characterized as a mutant that affects the normal accumulation of cuticular lipids on Arabidopsis stems (Koornneef et al., 1989; Mcnevin et al., 1991; Jenks et al., 1995). The subsequent molecular characterization of this locus established the sequence homology between *GL2* and *CER2* (Negruk et al., 1996; Xia et al., 1996), and these two genes proved to encode archetypal members of the BAHD family proteins (D’Auria, 2006). More recently we compared the functionality of the *Gl2* and *Gl2-like* genes by genetic complementation assays, relative to the function of the Arabidopsis *CER2* gene (Alexander et al., 2020).

In this study we illustrate the application of two specific analytical capabilities that overcome shortcomings associated with traditional “bulk” tissue characterizations. Moreover, many such reported studies of cuticular lipid mutants are limited to assays that measure metabolic outcomes taken at a single time-point, and thus ignore the dynamics of the system.

Furthermore, the use of more assiduous analytical technologies enabled the detection and measurement of cuticular lipid accumulation patterns in discreet tissue samples. Specifically, we used an on-column large volume injection capabilities of the LVI-PTV injector (Engewald et al., 1999; Wilson et al., 2000) that concentrates dilute samples during the gas chromatography (GC)-injection process, and thus assayed the dynamics of cuticular lipid accumulation during the development of single Arabidopsis flowers. We coupled this capability with the use of MALDI-MSI (Cha et al., 2009; Feenstra et al., 2017; Jun et al., 2010; Lee et al., 2012), to examine the changes in the distribution of cuticular lipids on developing Arabidopsis wild-type and *cer2* mutant flower surfaces, and obtained dynamic cuticular lipid data, at spatial resolution of a few cells.

## RESULTS

### Developing Flowers as an Informative System on the Dynamics of Extracellular Cuticular Lipid Deposition

The relatively rapid rate of tissue differentiation and development that occurs during the emergence of Arabidopsis flowers provides a convenient system to explore the dynamics of extracellular cuticular lipid accumulation. In addition, by using the LVI-PTV injector capabilities (Engewald et al., 1999; Wilson et al., 1997) to concentrate dilute samples during the GC-injection process, we characterized the extracellular cuticular lipid profiles extracted from individual flowers, collected at different stages of flower development. These attributes enabled the dynamic comparison of the cuticular lipid accumulation during flower development of wild-type and *cer2-5* mutant plants, and the effect of genetically complementing this mutation by the transgenic expression of a wild-type *CER2* allele.

The developmental stages of flowers were defined from their morphological appearances, as follows (Figure 1): Stage A, are flowers with closed buds; Stage B flowers are identified by the first emergence of petals from the bud; at Stage C the emerging petals are perpendicular to the flower axis; and at Stage D the flower recloses and the developing silique begins to emerge. Time lapse videos of 2 to 4 replicate flowers enabled the timing of these 4 stages, relative to Stage A, which was assigned the zero-time point. Thus, flowers on wild-type plants reached Stage B at 2.10 + 0.05 h, Stage C occurred at 9.50 + 0.96 h, and Stage D occurred at 33.5 + 2.6 h. Statistical analysis of these data show that the B to C transition is slowed by the *cer2-5* mutation, but this delay is alleviated by the transgenic expression of *CER2* in this mutant (Figure 2; Supplemental Table 1). In parallel, we determined the biomass of these flowers as a quantitative measure of flower growth (Figure 3; Supplemental Table 2). Thus, although the *cer2-5* mutation affects the developmental timing of the flower, it does not alter the biomass of the resulting flowers (Figure 3).

**Figure 1.**
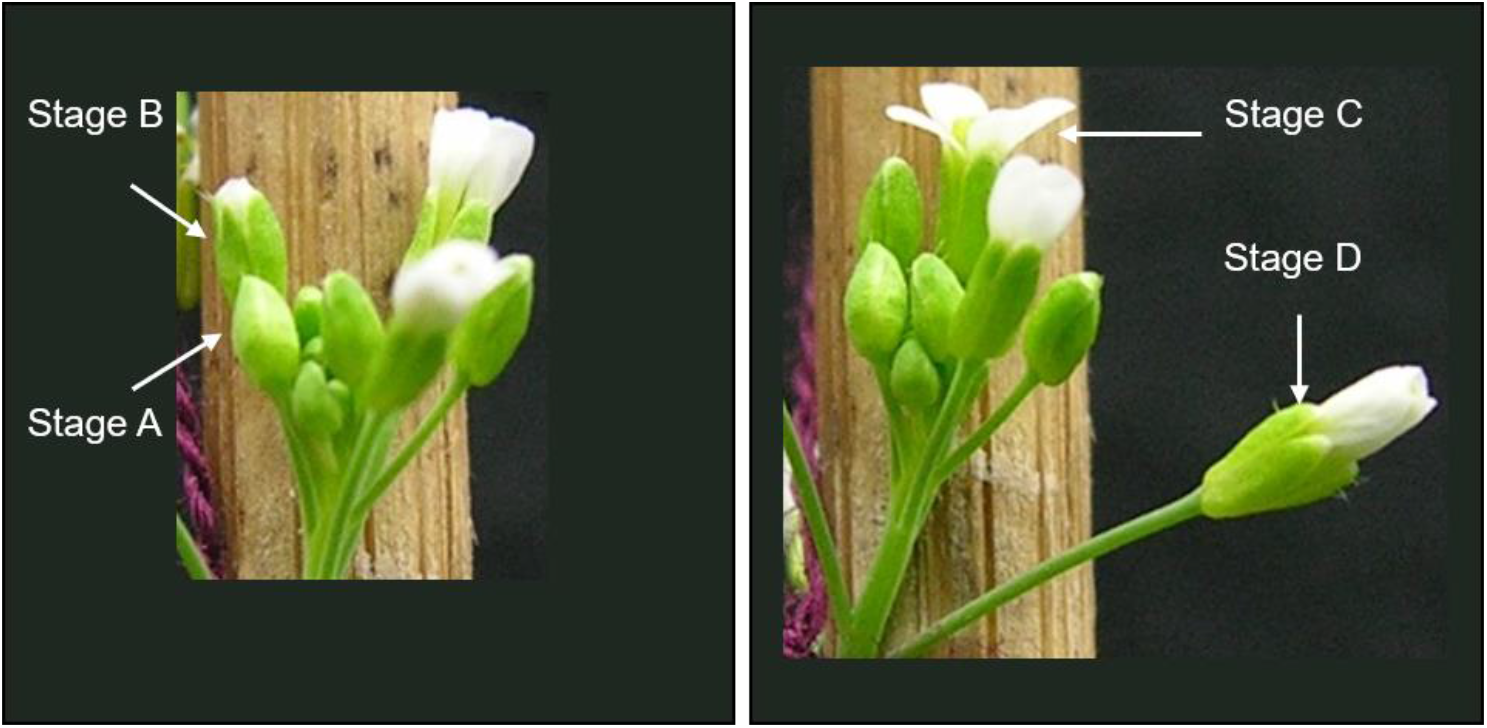
Arabidopsis flower development. Stage A are flowers with closed buds; Stage B flowers are identified by the first emergence of petals from the bud; Stage C, the emerging petals are perpendicular to the flower axis, and at Stage D the flower recloses and the developing silique begins to emerge.

**Figure 2.**
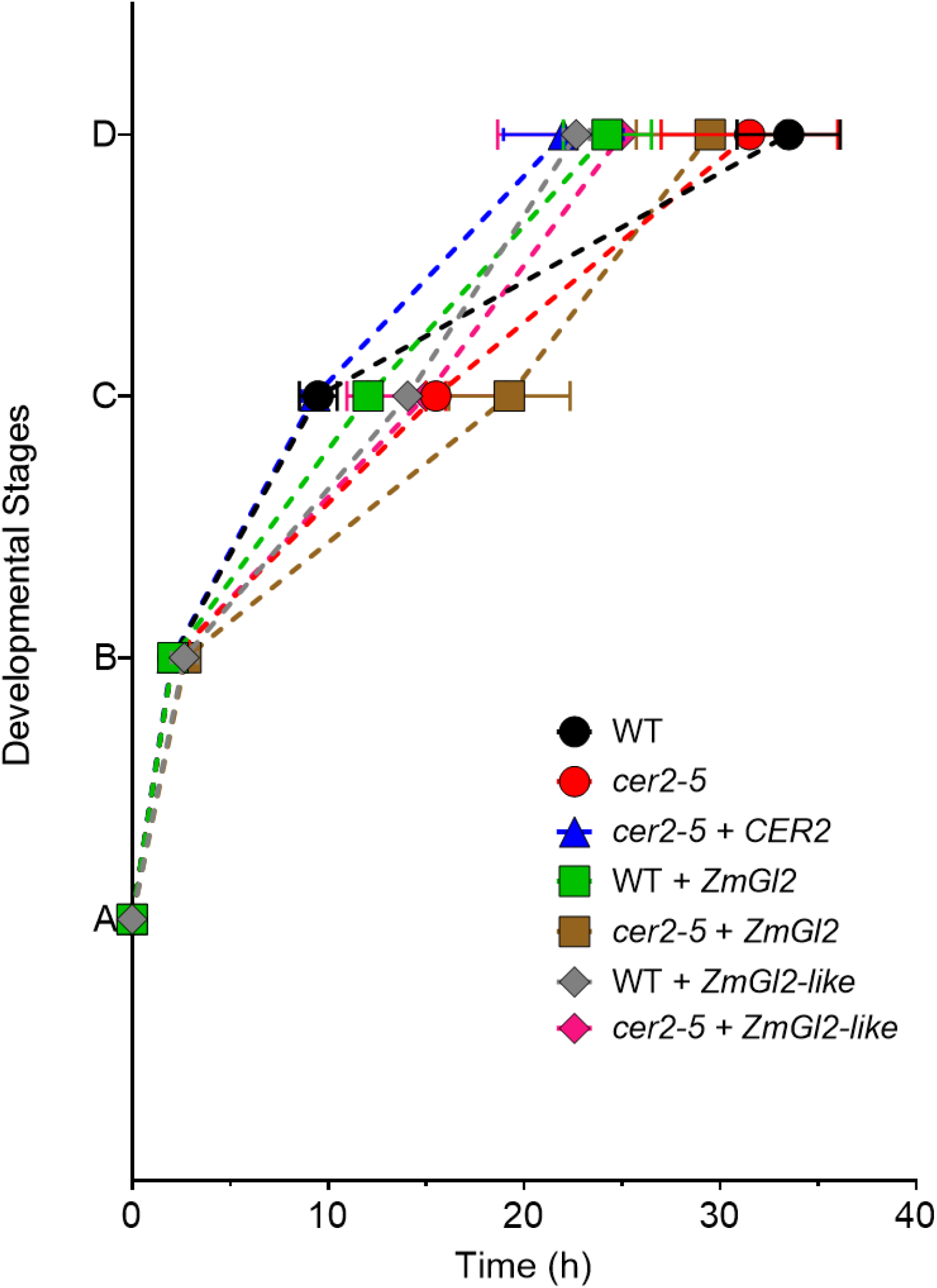
Timing of Arabidopsis flower development. Timing of transitions from Stage A to Stages, B, C, and D of non-transgenic and transgenic wild-type and *cer2-5* mutant flowers expressing either the *CER2* gene or the maize *Gl2* or *Gl2-like* genes. The data is the average of 2-4 replicates + standard error, determined from time-lapse videos.

**Figure 3.**
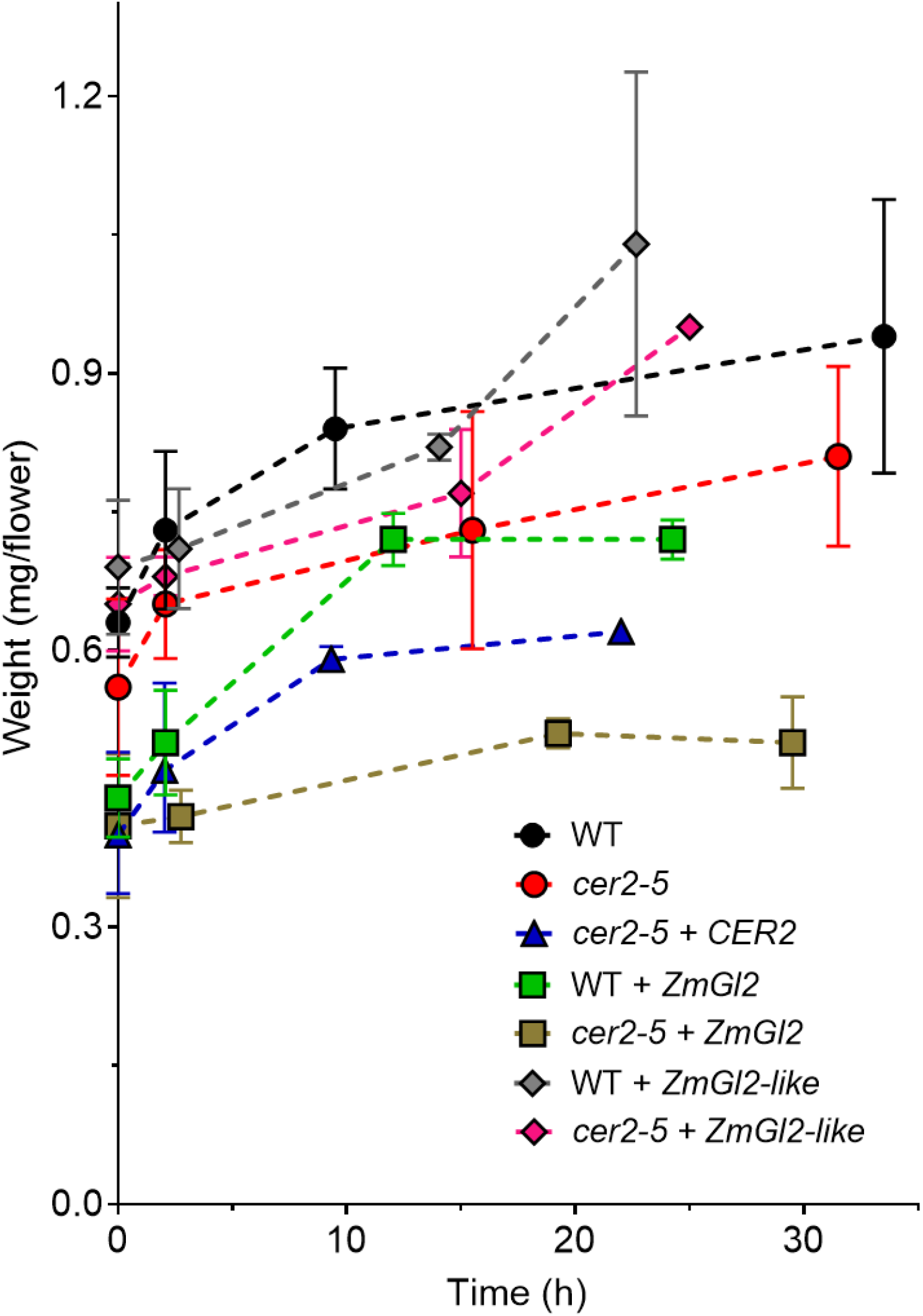
Change in biomass with flower development. Fresh weight of flowers + standard deviation is plotted against time taken for Arabidopsis flowers over-expressing maize transgenes *Gl2* or *Gl2-like* to develop. Biomass of Arabidopsis flowers. The fresh weight of 6-13 pooled flowers of the indicated genotypes were weighed at each of the 4 flower stages defined in Figures 1 and 2. The data is the average of 2-3 determinations + standard deviation.

Analysis of the extracellular cuticular lipid load on these developing flowers reveals dynamic changes in the accumulation of the alkyl metabolites. These metabolites are fatty acids and derivatives, such as primary alcohols, alkanes (linear and iso-branched), ketones, and secondary alcohols (Supplemental Data Sheet 1). The accumulation of these lipids increases asymptotically with development, through an initial rapid rate of accumulation (i.e., the transition from Stages A to B) that is as high as 26.3 nmol of lipid/h/g of fresh weight, which slows to a rate of 1.5-2.6 nmol of lipid/h/g of fresh weight between Stages C to D (Supplemental Figures 1 and 2; Supplemental Tables 3 and 4). The effect of the *cer2-5* mutation is to nearly eliminate the initial burst in cuticular lipid accumulation and to reduce the later accumulation rate to near zero (Supplemental Figures 1 and 2; Supplemental Tables 3 and 4). As would be expected, the transgenic expression of *CER2* in the *cer2-5* mutant increases the rate of floral cuticular lipid accumulation to that found with wild-type flowers.

In the context of these genetic-based changes in the dynamics of the total cuticular lipid loads on the flowers, the major components are always hydrocarbons, which account for 70 to 80% of the quantified cuticular lipids (Figures 4–6; Supplemental Data Sheet 1). In the wild-type state, over 70-80% of the hydrocarbons are of 29-carbon chain length, with lesser amounts of other linear chains (25, 27 or 31 carbon atoms) or 2-methyl-branched chains (26, 28, 29, or 30 carbon atoms). The other less abundant classes of cuticular lipids, in descending order of abundance, are secondary alcohols, ketones, primary alcohols and fatty acids. The dynamic accumulation patterns of the majority of the individual metabolites follow a similar asymptotic curve as the total cuticular lipid accumulation pattern (Figures 4–6). Moreover, with the exception of the VLCFAs, the most abundant chain length of each of these alkyl derivatives match those of the hydrocarbons (i.e., either the same number of carbon atoms or one carbon atom longer than the hydrocarbon), as would be predicted from the currently accepted model of cuticular lipid biosynthesis pathway, i.e., VLCFA elongation coupled to either a reductive or decarbonylative branched pathway (Bianchi et al., 1985; Post-Beittenmiller, 1996). The one exception is the chain-length distribution of the VLCFAs, which are dominated by 20 to 24 carbon chain length, whereas the other alkyl classes are dominated by the longer chain lengths (28-30 carbon atoms).

**Figure 4.**
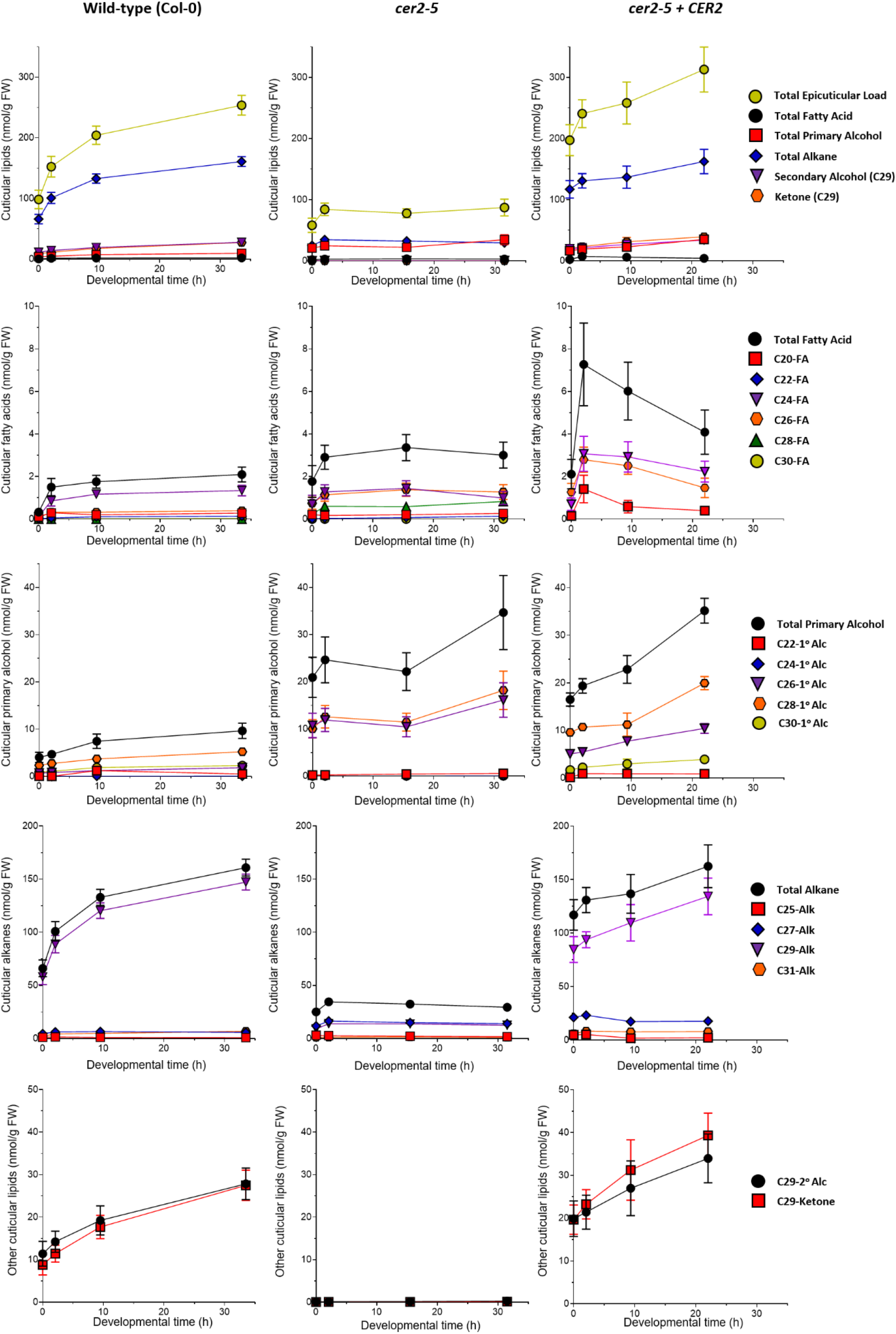
Cuticular lipid accumulation on developing Arabidopsis flowers. Cuticular lipids were extracted from individual Arabidopsis flowers of wild-type, *cer2-5* mutant, and *CER2* overexpressed in the mutant, and analyzed using a GC equipped with a LVI-PTV inlet injector (see Materials and Methods). The data represents the average + standard error, of two independent experiments for the wild-type and *cer2-5* mutant arranged in order of most abundant metabolites.

**Figure 5.**
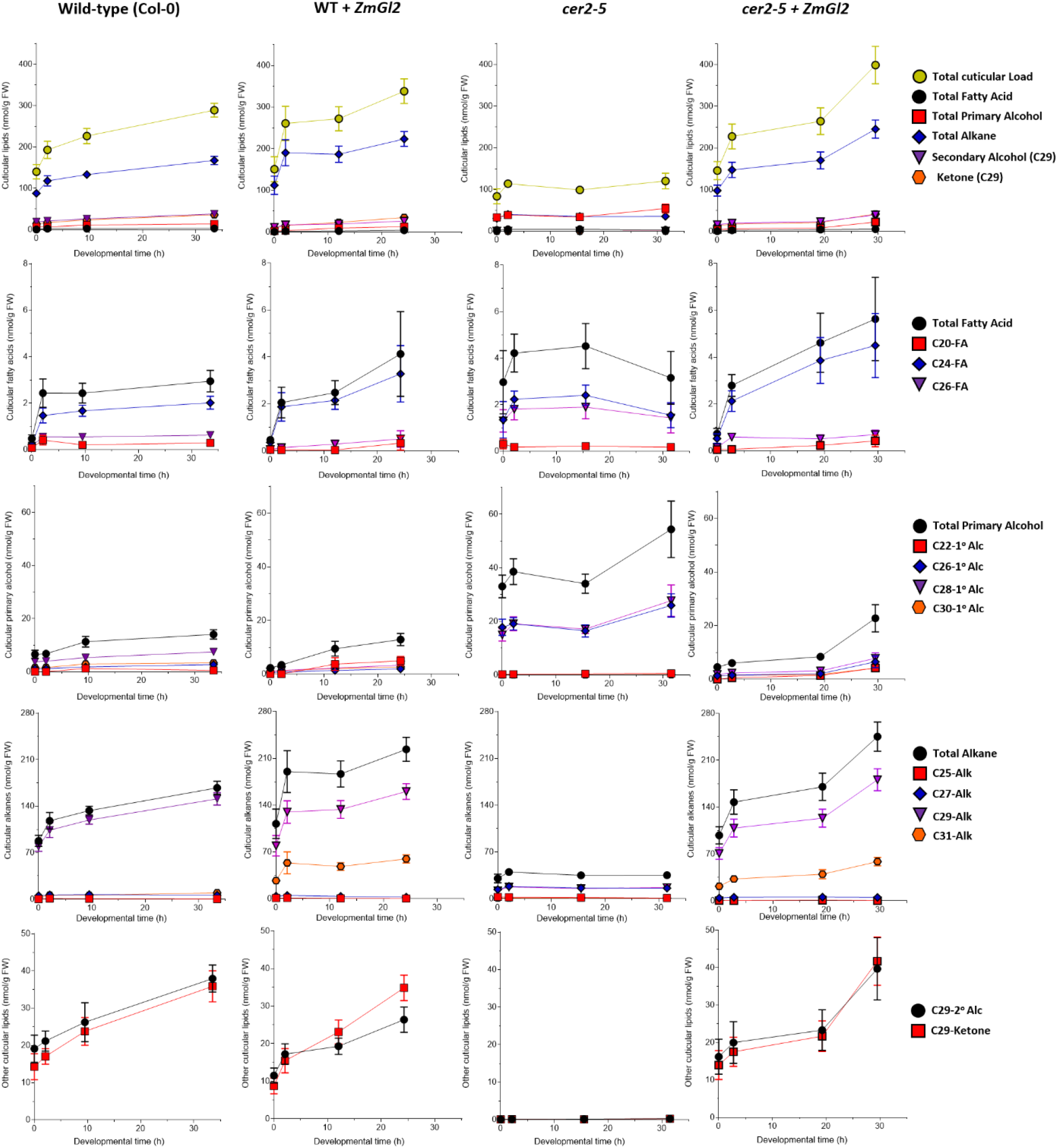
Cuticular surface lipid trends on developing Arabidopsis flowers. Cuticular lipids were extracted from individual flowers of genotypes: wild-type (Col-0), *cer2-5* mutant, and either of these lines expressing maize *Gl2* transgene. The data represent average + standard error of 6 replicates arranged in order of highest abundance.

**Figure 6.**
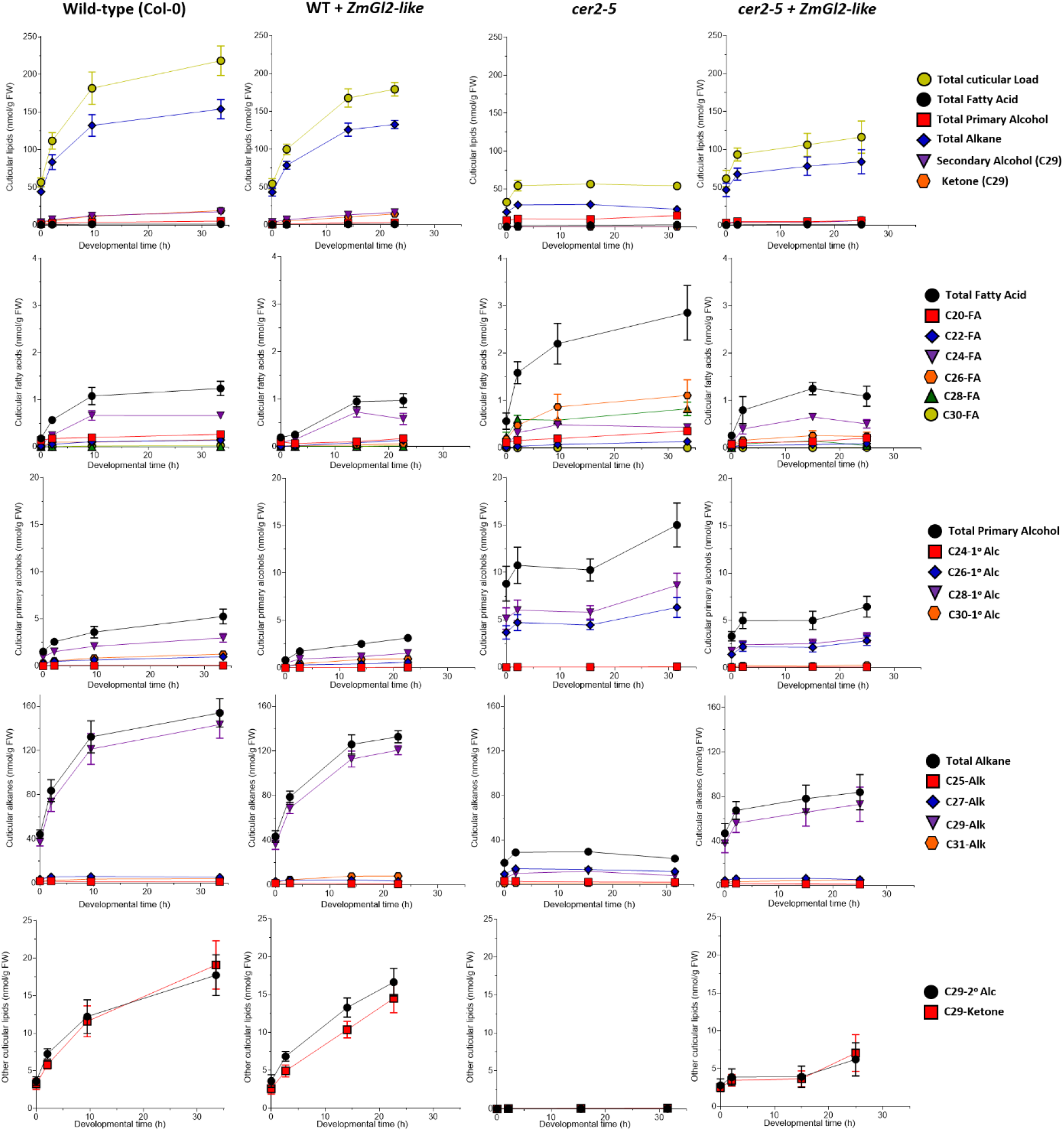
Cuticular lipid trends on developing Arabidopsis flowers. Cuticular lipids were extracted from individual flowers of genotypes: wild-type (Col-0), *cer2-5* mutant, and either of these lines expressing maize *Gl2-like* transgene. The data represent average + standard error of 6 replicates arranged in order of highest abundance.

Supplemental Tables 3 and 4 tabulates the rates of accumulation for each of the cuticular lipid metabolites between each of the four flower developmental transitions. While the asymptotic rates of accumulation are maintained for most individual components, in the *cer2-5* mutation at the initial stages of development (Stage A-B transition) these rates are approximately ½ the rate determined in the wild-type plants. Two exceptions to this generalization are the net accumulation rates of 26:0 and 28:0 fatty acids and primary alcohols of these chain lengths. The initial rate of accumulation of these metabolites is increased in the *cer2-5* mutant by 2-to 5-fold as compared to the wild type, and this increase is reversed by the transgenic expression of *CER2* (Figure 4; Supplemental Table 4).

### Transgenic Expression of the Maize *Gl2* and *Gl2-like* in the *cer2* mutant Restores Cuticular Lipid Accumulation

Using the above data as a baseline, two independent transgenic experiments were conducted to characterize the effect of expressing either the maize GL2 or GL2-LIKE proteins in a wild-type or *cer2-5* mutant background. The expression of either *Gl2* or *Gl2-like* in the *cer2-5* mutant does not affect the developmental timing of the flowers, however in the wild-type background both transgenes induce a faster rate of flower development (Figure 2 and Supplemental Table 1). Additionally, the transgenic expression of *Gl2* in both these genetic backgrounds reduced the mass of the flowers (Figure 3, and Supplemental Table 2), whereas the *Gl2-like* transgene did not impact the mass of the resulting flowers (Figure 3, and Supplemental Table 2).

The transgenic expression of either *Gl2* or *Gl2-like* in the *cer2-5* mutant affected an increase in the level of cuticular lipid accumulation on developing flowers. The *Gl2* transgene appears to be more effective in returning the accumulation of cuticular lipids to wild-type levels (Supplemental Figure 1), whereas the *Gl2-like* transgene only partially rescued this trait (Supplemental Figure 2). This latter effect is particularly noticeable in the accumulation pattern of the major components, the alkanes, ketones and secondary alcohols (Figure 6). Thus, whereas the transgenic expression of *Gl2* in the *cer2-5* mutant restores the levels of these lipids to wildtype levels, the transgenic expression of *Gl2-like* in this mutant leads to the accumulation of these lipids at only 50% of the levels found in the wild-type (Figures 5 and 6).

### Non-uniform Tissue and Cellular Level Spatial Distribution of Epicuticular Surface Lipids on the Abaxial and Adaxial Surfaces of Arabidopsis Flowers

The extraction-based data presented in Figures 4–6 and Supplemental Figures 1 and 2, integrates the cuticular lipid compositional changes that occur on a number of different flower tissues (e.g., petals, sepals, anthers and stigmatic tissues), each of which is on a different developmental schedule (Alvarez-buylla et al., 2010; Smyth, 1990). Therefore, we applied *in situ* mass spectrometric imaging, specifically MALDI-MSI technology (Lee et al., 2012; Kaspar et al., 2011; Dong et al., 2016), to individually localize these metabolites at a high spatial scale (~80 μm), which enables the visualization of the accumulation of these metabolites among the different tissues of the flower. Given that Arabidopsis epidermal cells present surface areas that are approximately 1000 to 4000 μm^2^ (Cookson et al., 2006; Kheibarshekan Asl et al., 2011; Robinson et al., 2018), and the laser spot size used in the MSI experiments has diameter of ~80 μm (i.e., 5000 μm^2^ area) we estimate that these MSI data are visualizing the surface distribution of cuticular lipids at a resolution as small as 2-4 cells. Moreover, MALDI-MSI was used to investigate the cuticular surface lipid distribution on both surfaces of the developing flowers (Figure 7A), which enabled the comparison of the distribution of individual metabolites on the adaxial (inner facing) and abaxial (outer facing) surfaces of sepals and petals, and the surfaces of anthers and stigma (Figure 7B).

**Figure 7.**
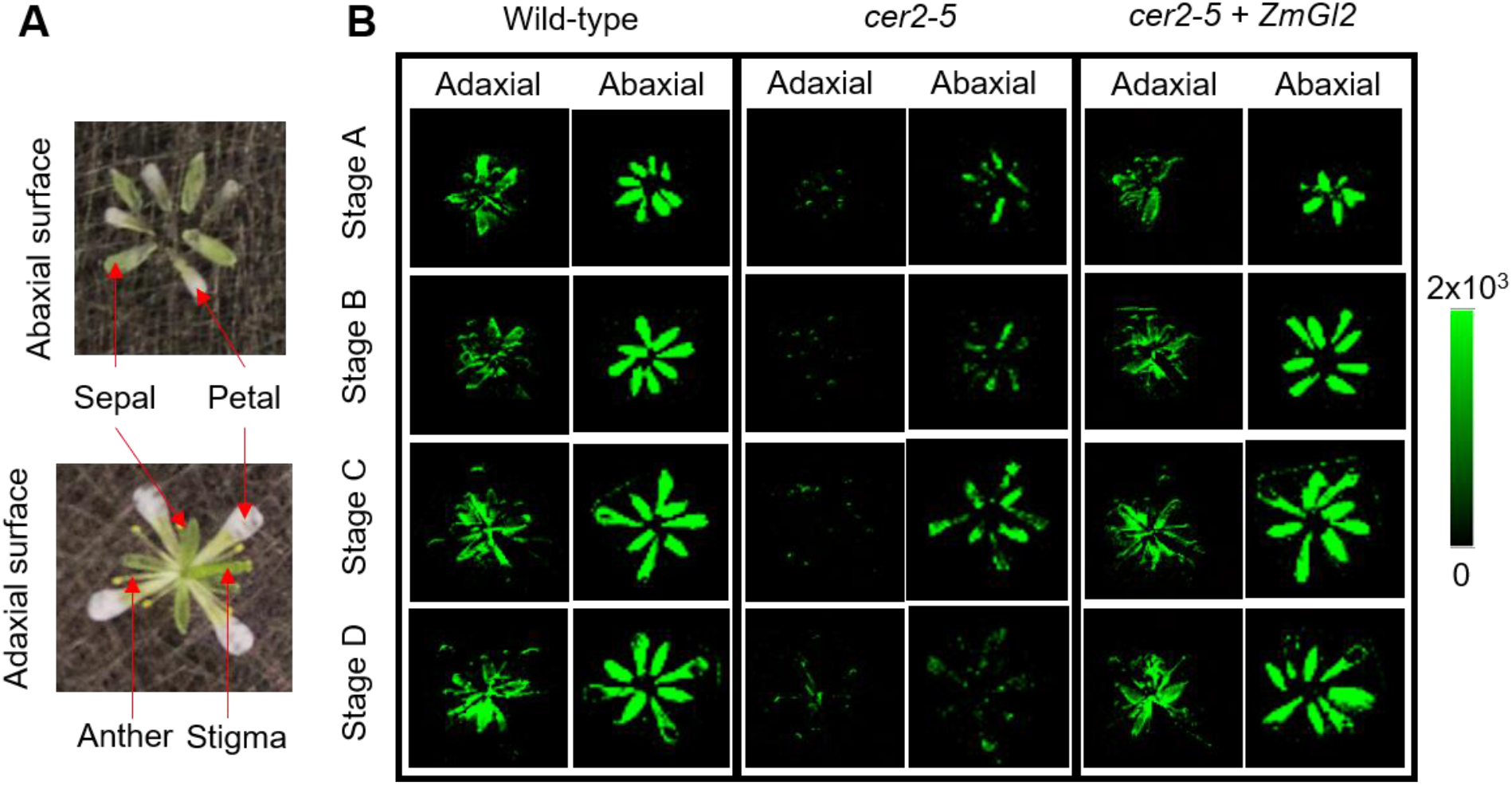
Spatial distribution of nonacosane on Arabidopsis flowers. A) Visual images of the adaxial and abaxial surfaces of Arabidopsis flowers positioned on a MALDI plate ready for mass spectrometric imaging. B) The distribution of nonacosane (C29 alkane) on the adaxial and abaxial floral surfaces at four stages of development (Stages A to D) of the indicated genotypes. MSI absolute intensity scale bar is color coded: green is maximum signal and black is minimum signal.

This imaging technology was used to determine the distribution of individual cuticular lipid metabolites at all four developmental stages of wild-type Arabidopsis flowers, and these images were compared to flowers of the *cer2-5* mutant, and to the *cer2-5* mutant that was transgenically expressing the *Gl2* gene. Figure 7B shows representative data revealing the surface distribution of the most abundant cuticular lipid (C29 alkane), which is evenly distributed on both the adaxial and abaxial floral surfaces of wild-type flowers during the later stages of development (Stages C and D). However, at the earlier stages of foliar development (Stages A and B), this cuticular lipid is primarily detected on the abaxial surface and occurs at reduced levels on the adaxial surface of the flower (Figure 7B). The *cer2-5* mutation significantly reduced the amount of the C29 alkane on both the adaxial and abaxial floral surfaces, and the transgenic expression of the maize *Gl2* in this mutant background reversed this effect, so that the distribution of this lipid is similar to the distribution found on the wild-type flowers.

Because Stage C flowers display the clearest physical separation of the different flower tissues, MSI analysis was used to evaluate the distribution of a wider range of cuticle metabolites among flower tissues in response to the transgenic expression of *Gl2* and *Gl2-like* in the *cer2-5* mutant background (Figure 8). These imaging experiments specifically focused on the spatial distribution of cuticle metabolites that are metabolically derived from the 30:0 fatty acid, namely C30 aldehyde, and the C29 alkane, secondary alcohol and ketone. These images reveal that whereas these metabolites are evenly distributed on the abaxial surfaces, they more discreetly localized to the petals, and edges of the sepals of the adaxial surface of wild-type flowers. The *cer2-5* mutant flowers show uniform reduction in the levels of C30 aldehyde, C29 alkane and C29 ketone on both surfaces of the flower sepals and petals, and the 30:0 fatty acid and the C29 secondary alcohol are at undetectable levels. The transgenic expression of *Gl2* or *Gl2-like* in the *cer2-5* mutant not only increased the levels of these metabolites, but also reiterated the distribution patterns of these metabolites to that imaged on the wild-type flowers (Figures 8A and 8B).

**Figure 8.**
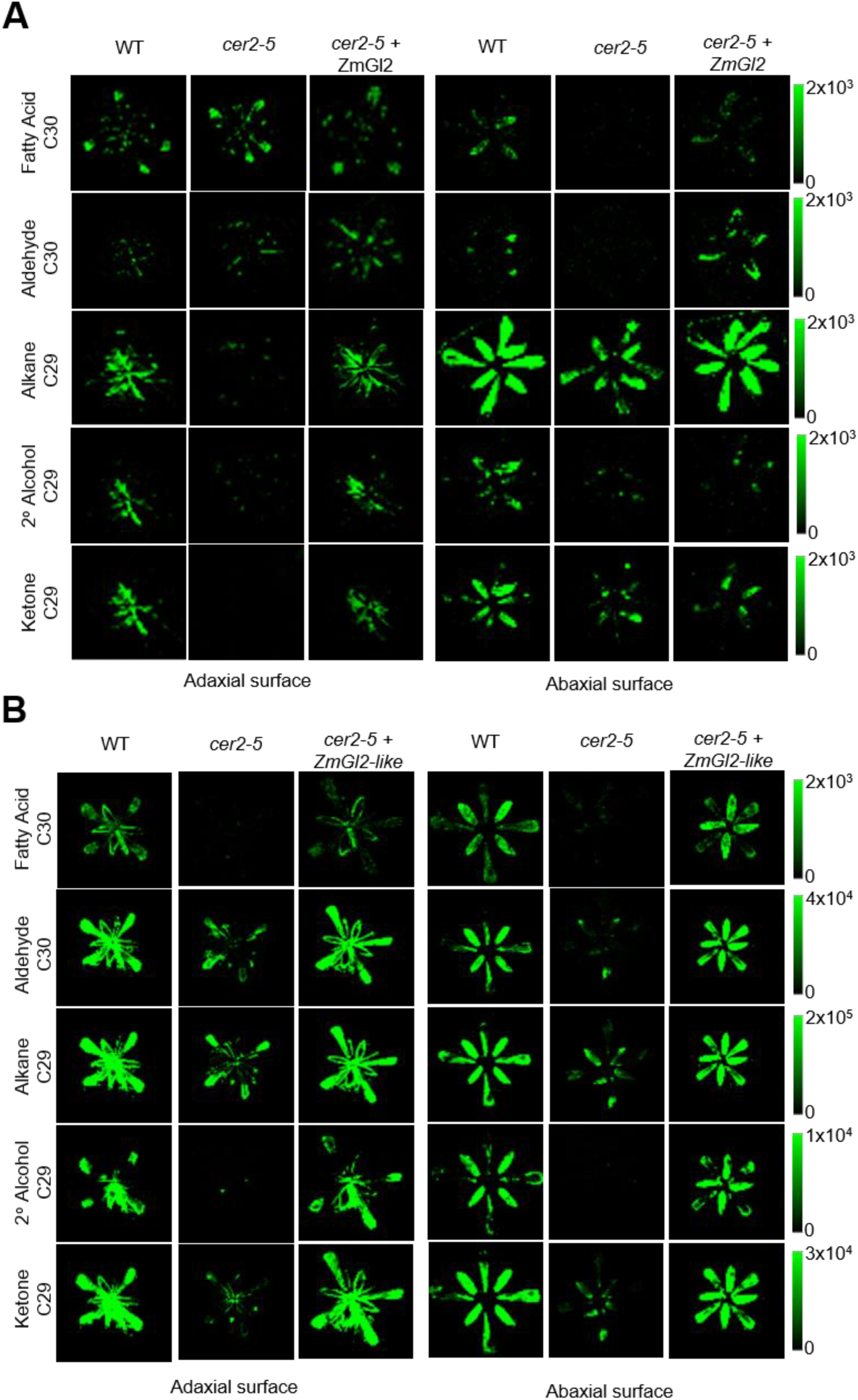
Spatial distribution of cuticular lipids on the adaxial and abaxial surfaces of Arabidopsis flowers. Two independent experiments that visualize the effect of the transgenic expression of *Gl2* (A) or *Gl2-like* (B) on the spatial distribution of 30:0fatty acid and alkyl-derivatives on the two surfaces of Arabidopsis flowers. The genotype of each flower is indicated and MSI absolute intensity scale bar is color coded: green is maximum signal and black is minimum signal.

### MSI can be used to Visualize the Spatial Distribution of Individual Metabolic Conversions of the Cuticular Biosynthetic Network

An additional application of MSI technology is the ability to infer the spatial distribution of the underlying metabolic conversion processes that support a metabolic network. We illustrate this by the visualization of the co-localization of substrate-product pairs of individual metabolic reactions of the cuticular biosynthetic pathway. For example, consider the distribution of pairs of fatty acids, 24:0, 26:0, 28:0, 30:0 and 32:0; each sequential pair has a precursor-product relationship in the reactions of fatty acid elongation. Additionally, each fatty acid has a precursor-product relationship with an aldehyde of the same carbon chain length. Figures 9 and 10 and Supplemental Figures 3 and 4 visualize the spatial distribution of these precursor-product relationships by false color coding of each substrate (green) and each product (red) of these metabolic conversions. In the resulting fused images, the yellow colored zones indicate the spatial co-location of these precursor-product pairs, indicative of the underlying tissue/cell locations where this metabolic conversion may be occurring.

**Figure 9.**
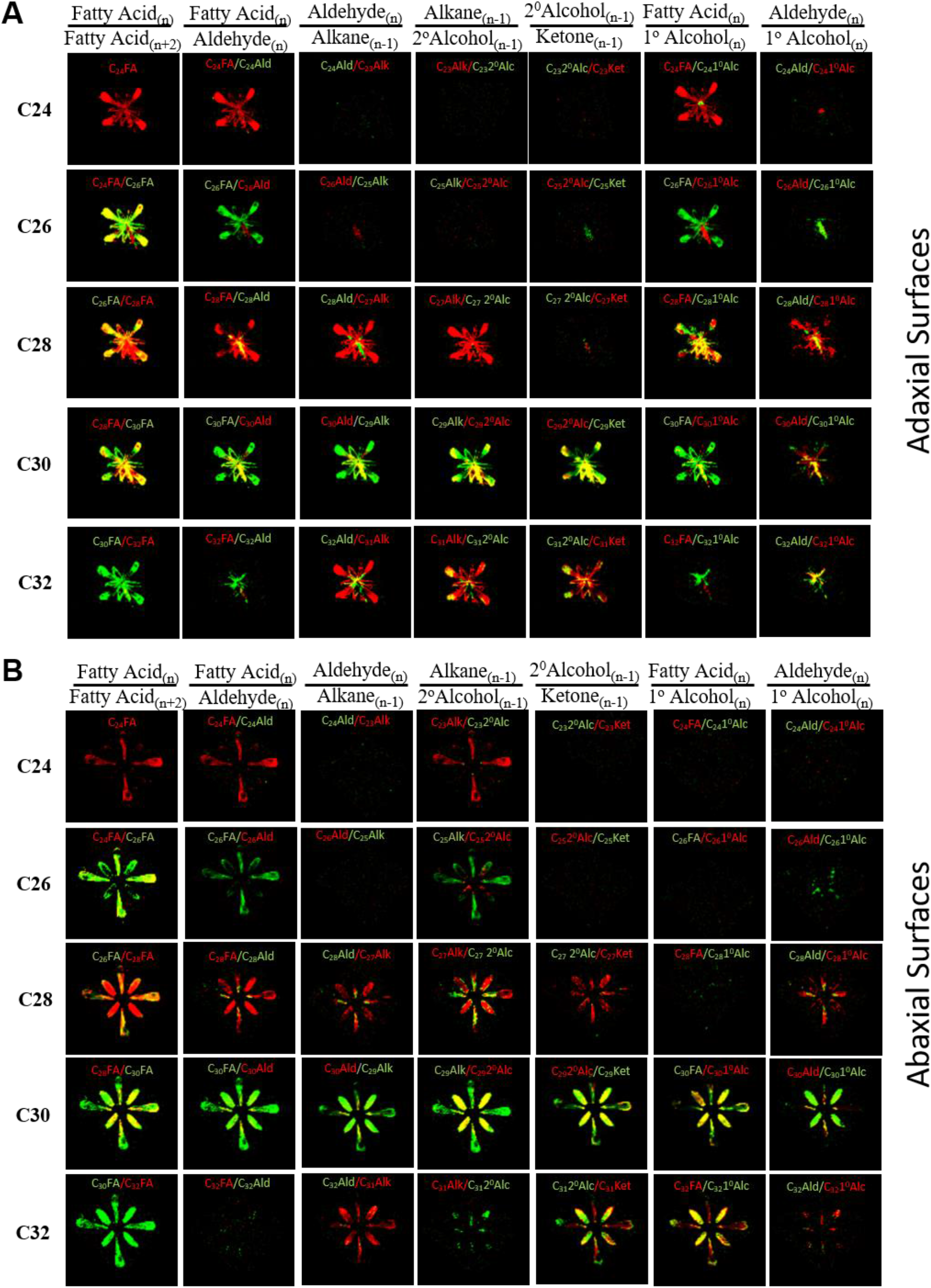
Mass spectrometric imaging of the co-localization of substrate-product pairs of individual reactions of the cuticular lipid biosynthesis pathway. Each image is a fusion of two false color-coded MSI images, green for the substrate, and red for the product of a single metabolic reaction that contributes to cuticular lipid biosynthesis pathway. In these fused images, yellow represents the spatial zone where these substrate-product pairs co-localize, indicating the location of the underlying tissue/cells where this metabolic interconversion could be occurring. The distribution of 24:0 to 32:0 fatty acid and their alkyl derivatives on the adaxial (A) and abaxial (B) floral surfaces wild-type Col-0 flowers.

**Figure 10.**
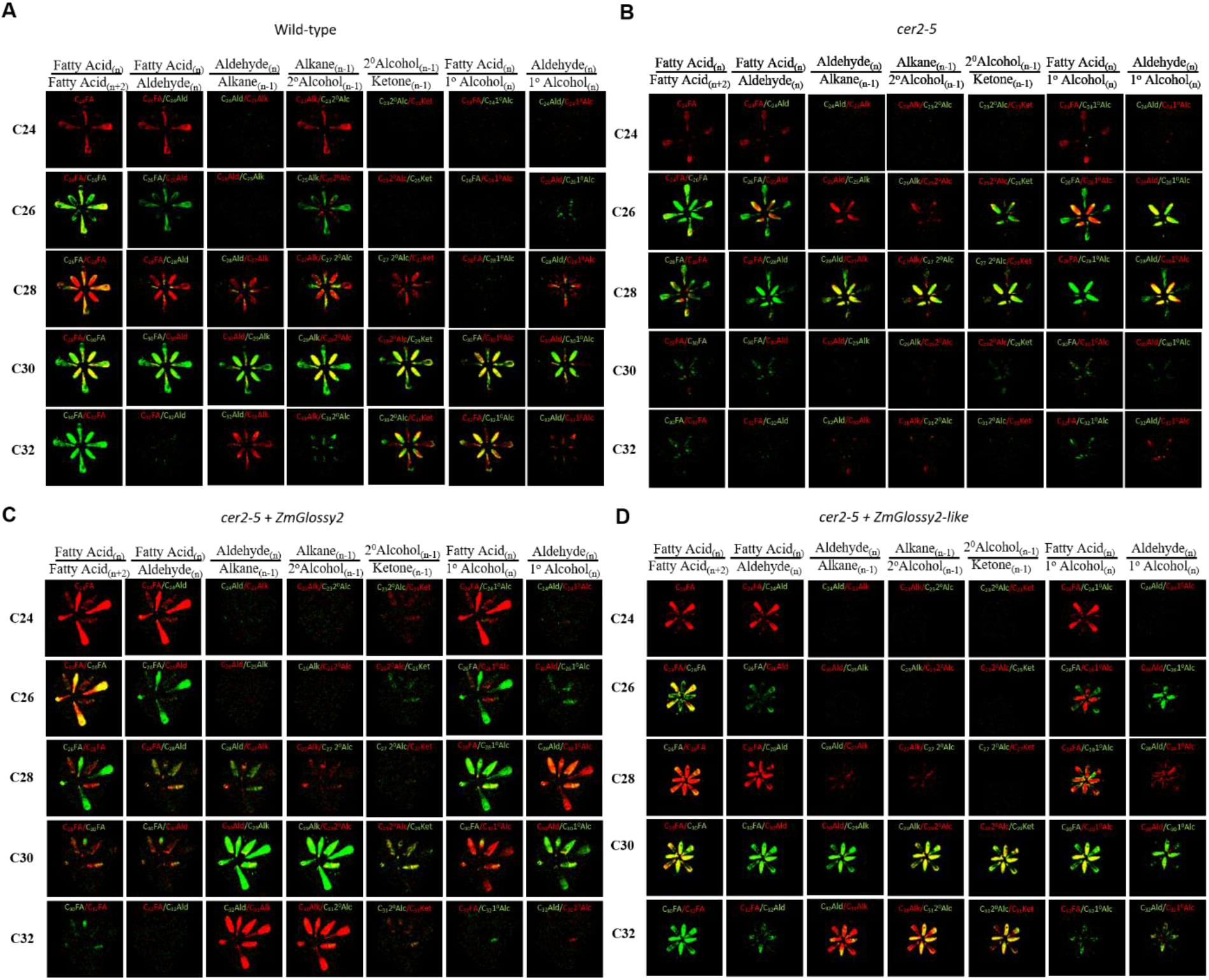
Mass spectrometric imaging of the co-localization of substrate-product pairs of individual reactions of the cuticular lipid biosynthesis pathway. Each image is a fusion of two false color-coded MSI images, green for the substrate, and red for the product of a single metabolic reaction that contributes to cuticular lipid biosynthesis pathway. In these fused images, yellow represents the spatial zone where these substrate-product pairs co-localize, indicating the location of the underlying tissue/cells where this metabolic interconversion could be occurring. The distribution of 24:0 to 32:0 fatty acid and their alkyl derivatives on the abaxial floral surfaces are indicated for wild-type Col-0 (A), *cer2* mutant (B), and *cer2* mutant transgenically expressing *Glossy2* (C) or *Glossy2-like* (D).

Figure 9A shows that the fatty acid precursor-product pairs of the fatty acid elongation reactions predominately co-localize on the adaxial surface of wild-type flower petals. In contrast, there is less such co-localization on the abaxial surface (Figure 9B). Similar co-localization occurs on the adaxial surfaces of the stigmatic tissue and petals (Figure 9A) for the four reactions that lead from 30:0 fatty acid to the C29 ketone (i.e., C_30_ FA-C_30_ Ald, C_30_ Ald-C_29_ Alk, C_29_ Alk-C_29_ 2^0^Alc and C_29_ 2^0^Alc-C_29_ Ket). And such co-localization is found on the abaxial surface of sepals for the last two reactions of decarbonylative branch (i.e., C_29_ Alk-C_29_ 2^0^Alc and C_29_ 2^0^Alc-C_29_ Ket) (Figure 9B). This contrasts with the substrate-product pairs of the reductive branch of the pathway that generates the primary alcohols (i.e., C_28_ FA-C_28_ 1^0^Alc and C_30_ FA-C_30_ 1^0^Alc pairs), these co-localize on the abaxial surface of sepals and stigmatic surfaces (Figure 9).

The *cer2-5* mutation does not affect these co-localization patterns, with the exception of products that are generated from 28:0 and 30:0 fatty acids. Thus, consistent with the proposed role of CER2 in affecting the 28:0 to 30:0 fatty acid elongation process (Hannoufa et al., 1993; Jenks et al., 1995), the 28:0-30:0 fatty acid substrate-product pairs do not co-localize in the *cer2-5* mutant (Figure 10B, Supplemental Figure 3A). As a result, there are altered spatial accumulation patterns of the C28 alkyl derivatives within both the decarbonylative and reductive branches of the cuticular lipid pathway. Specifically, in the *cer2-5* mutant the substrate-product pairs from C28 aldehyde to C27 ketone primarily co-localize in the sepals, and the C26 aldehyde-C26 primary alcohol and C28 aldehyde-C28 primary alcohol substrate-product pairs primarily co-localize in the sepals and stigmatic tissue surface. These differential co-localizations in the *cer2-5* mutant may indicate that both the decarbonylative and reductive branches of the cuticular lipid pathway occur in sepal tissues, but the reductive branch of the pathway appears to predominate in the stigmatic tissues. The transgenic expression of *Gl2-like* in the *cer2-5* mutant background restores the co-localization pattern of the 28:0 and 30:0 fatty acid elongation substrate-product pairs and the decarbonylative and reductive pathway substrate-product pairs to the pattern observed in the wild-type flowers (Figure 10D, Supplemental Figure 3B). Such a restoration of the substrate-product patterns is also obtained when the *Gl2* is transgenically expressed in the *cer2* mutant (Figure 10C).

These substrate-product co-localization patterns were also identified and compared at different stages of flower development (Supplemental Figure 4, Supplemental Figure 5). These data are presented only for the abaxial surfaces of the flowers because the signal-to-noise ratios obtained from these surfaces were sufficiently high to enable the formulation of robust conclusions (Supplemental Figure 4), whereas this was not the case for the data gathered from the adaxial surface (Supplemental Figure 5). During the development of wild-type flowers, the area of co-localization of the fatty acid elongation substrate-product pairs (24:0 to 28:0) that occurs in the petals, decrease as the flowers proceed to Stage D, whereas the co-localization of the 26:0 to 30:0 fatty acid elongation substrate-product pairs becomes concentrated to the sepals (Supplemental Figure 4). The *cer2-5* mutation disrupts these co-localization patterns, so that throughout flower development only the 24:0-26:0 fatty acid pair co-localize in petals, whereas, the 26:0-28:0 fatty acid pair co-localize to the sepals (Supplemental Figure 4). The transgenic expression of *Gl2* in the *cer2-5* mutant did not affect these altered co-localization patterns of the 24:0-26:0 fatty acid substrate-product pairs, and no co-localization patterns were observed for the 26:0-28:0 fatty acid pair (Stages C and D; Supplemental Figure 4).

In wild-type flowers the spatial pattern of the 28:0-30:0 fatty acid elongation substrate-product pairs are distinct from the substrate-product pairs of the decarbonylative and reductive branches of the pathway that initiate from these two fatty acids. Namely, the C28 and C30 aldehydes and alcohols, and the C27 and C29 derived alkanes, ketones and secondary alcohols co-localize to the abaxial surfaces of sepals, and this pattern is most apparent at the early stages of flower development (Stages A and B; Supplemental Figure 4). However, later in flower development these patterns become less co-localized (Stages C and D; Supplemental Figure 4). In contrast, in the *cer2-5* flowers these co-localization patterns are altered, particularly for the substrate-product pairs that lead from C25 and C27 alkanes to the analogous ketones, and the substrate-product pairs C26 aldehye-C26 primary alcohol. Specifically, the sepal co-localization patterns for C25 alkane to C25 ketone pairs are maintained only during the early stages of flower development (Stages A and B; Supplemental Figure 4), whereas the C27 alkane to C27 ketone pairs, and C26 aldehyde-primary alcohol pair are maintained throughout development (Supplemental Figure 4). These highlighted *cer2*-dependent alterations in the co-localization of substrate-product pairs are consistent with the proposed role of CER2 in affecting the 28:0 to 30:0 fatty acid conversion resulting in the higher accumulation of precursor fatty acid products, namely 26:0 and 28:0 (Hannoufa et al., 1993; Jenks et al., 1995; Haslam et al., 2012). The transgenic expression of *Gl2* in the *cer2-5* mutant background restored co-localization of the substrate-product pairs patterns for the C30 alkyl decarbonylative and reductive pathway on sepals, at least at the early stages of flower development (Stages A and B; Supplemental Figures 4), but not at the later stages of development (Stages C and D; Supplemental Figures 4). Therefore, the ability of the *Gl2* transgene to restore the spatial patterns of the C28 and C30 alkyl chain substrate-product pairs to near wild-type patterns, which were disrupted by the *cer2* mutation, further supports recent studies that indicated the functional homology between *Gl2* and *CER2* (Alexander et al., 2020).

## DISCUSSION

Elucidating the biochemical mechanisms regulating plant metabolic processes are complicated by the multicellular nature of these organisms. Specifically, this complexity is the consequence of the fact that molecular machineries can be distributed among different subcellular and/or cellular compartments, which are coordinated to achieve net metabolic interconversions (De Luca and St Pierre, 2000; Krueger et al., 2011; Lunn, 2007). This complexity is particularly associated with deciphering the extracellular cuticular lipid biosynthetic pathway that is restricted to the epidermal cells of aerial plant organs, and the products of this pathway are unidirectionally transported from these cells to the outer surface of plant organs. Experimental data from molecular genetic characterizations of *gl* and *cer* mutants that affect normal deposition of cuticular lipids are often interpreted relative to a core cuticular lipid biosynthesis pathway, which appear to be common to both maize and Arabidopsis, if not all plants (Bianchi et al., 1978; Bianchi et al., 1985; Kolattukudy, 1970; Kunst and Samuels, 2009; Lee and Suh, 2013; Post-Beittenmiller, 1996). This pathway integrates fatty acid elongation with either a reductive branch or a decarbonylative branch to generate a diverse set of alkyl lipid products. In this study we coupled genetic complementation with unique analytical strategies to measure and visualize distinct spatial and temporal patterns of cuticular lipid accumulation within floral tissues during flower development.

### Dynamics of the Distribution of Cuticular Surface Lipids

The majority of past biochemical and molecular studies of the accumulation of cuticular lipids have primarily been limited to a single time-point of analysis, overlooking the dynamics of the processes that integrate the developmental program of the tissue/organs that are being evaluated. Furthermore, most such data are collected from the analysis of cuticular lipid extracts prepared from relatively large biological samples that combine lipids generated by a heterogeneous collection of cells and tissues that are most probably of different age and different developmental states. In this study of developing flowers, the application of new, more assiduous analytical technologies provided the platform to address both of these potential limitations to decipher the physiology of cuticular lipid accumulation patterns.

Specifically, we investigated the effect of the *cer2* mutation and the transgenic expression of maize *Gl2* and *Gl2-like* transgenes on cuticular lipids of Arabidopsis flowers, monitoring the accumulation pattern of these lipids during the course of floral development. Initial characterization of *cer2* mutant have been interpreted to indicate that this gene product affects the elongation of fatty acids between 26:0 or 28:0 to 30:0 (Hannoufa et al., 1993; Jenks et al., 1995), a hypothesis that has been further elucidated by partial reconstitution experiments of the fatty acid elongation process in yeast (Haslam et al., 2012; Haslam et al., 2015).

In this study we gathered data with cuticular lipid extracts prepared from individual flowers at four different stages of development. In parallel, we also applied MALDI-MSI to image the spatial distribution of extracellular cuticular lipid metabolites on either the abaxial or adaxial surfaces of the developing flowers at a sufficiently high spatial resolution to distinguish between four different floral organs (petals, sepals, anthers and stigmatic surfaces) and determined the effect of each genetic manipulation on the extracellular cuticular lipid profiles.

These studies of the dynamics of flower development demonstrate that a block in a metabolic process that specifically occurs in the epidermal cells of the flower tissues (i.e., extracellular cuticular lipid deposition caused by the *cer2* mutation) does not affect the developmental timing of the integrated organ, but it does affect the biomass of the flower, particularly when the *cer2* mutation is complemented by the two maize homologs. The expression of the maize homologs of *CER2*, *Gl2-like* or *Gl2* transgenes in the *cer2-5* mutant reversed the effect of blocking the accumulation of cuticular lipids on flowers, but these transgenes have differential effects on the growth of the floral organs, where expression of *Gl2* reduced the flower biomass, whereas expression of *Gl2-like* does not alter the flower biomass. These findings indicate therefore, that the flower biomass trait can be affected by the deposition of the cuticle, possibly associated with the ability of the organ to retain water that is lost via transpiration through a “leaky” cuticle (Goodwin and Jenks, 2005).

During flower development the majority of the individual cuticular lipid components accumulate at near parallel, asymptotic patterns as the flower developed. The maximum observed rate of cuticular lipid accumulation occurs during the earlier stages of flower development, and this is at a rate of ~26 nmol/h/g fresh weight. This rate defines the minimal rate of *de novo* fatty acid biosynthesis that is required to feed the FAE system with stearoyl-CoA precursor. Furthermore, because the final products of the cuticular lipid biosynthetic pathway are components that are predominantly derivatives of 30-carbon fatty acids, 6 elongation cycles of the FAE system are required to elongate the 18:0 fatty acid precursor to these final products. Because each elongation cycle requires 1 molecule of malonyl-CoA generated from the cytosolic acetyl-CoA pool, and 2 molecules of NADPH, one can infer the minimal rates of acetyl-CoA (156 nmol/h/g fresh weight) and NADPH (312 nmol/h/g fresh weight) generation in the cytosol that is required to support the observed rate of cuticle lipid deposition. Prior characterizations of ATP-citrate lyase (Fatland et al., 2002; 2005) and the cytosolic homomeric acetyl-CoA carboxylase (Baud et al., 2003; Konishi and Sasaki, 1994; Yanai et al., 1995), which sequentially generate the cytosolic acetyl-CoA and malonyl-CoA pools, appears to indicate that there is sufficient activity of these enzyme in Arabidopsis tissue to provide this rate of acetyl-CoA and malonyl-CoA generation. Similarly, there appears to be sufficient activities of NADPH-generating enzymes (NADP+-dependent malic enzyme, glucose-6-phosphate dehydrogenase, and 6-phosphogluconate dehydrogenase) to support these needs of the FAE system (Wakao et al., 2008; Wakao and Benning, 2005; Wheeler et al., 2005; Yin and Ashihara, 2008)

The effect of the *cer2-5* mutation is to reduce the rate of cuticular lipid accumulation at the early stages of flower development by about 55% (to ~12 nmol/h/g fresh weight), and this rate plateaus to near zero later in flower development. As expected the transgenic expression of *CER2* in the *cer2-5* mutant background reverses the initial rate of cuticular lipid accumulation to that of wild-type state, enabling the normal levels of cuticular lipid accumulation. Similarly, the transgenic expression of *Gl2* (but not *Gl2-like*) also reverses the effect of the *cer2* mutation, increasing the rate of cuticular lipid accumulation to that of wild-type levels. In contrast, *Gl2-like* transgene increases the rate of cuticular lipid accumulation but does not fully restore this rate to the wild-type level.

In all these genetically complemented lines the near asymptotic pattern of cuticular lipid accumulation is shared by nearly all individual cuticular lipid components, and the rates of accumulation are higher than in the *cer2-5* mutant. The exception to this generalization is the accumulation pattern displayed by the primary alcohols and fatty acids, particularly of 26- and 28-carbon chain length, which increase in accumulation in the *cer2-5* mutant as compared to the wild-type flowers. The transgenic expression of *Gl2* or *Gl2-like* in the *cer2-5* mutant reverses these alterations in the accumulation of C26 and C28 primary alcohols and fatty acids (i.e., reduces accumulation) to the wild-type state. In combination, these genetic complementation experiments reinforce the conclusion that 2 maize genes have overlapping functionalities that are homologous to the *CER2* gene function (Alexander et al., 2020). Furthermore, these data may suggest the existence of two parallel fatty acid elongation processes, one feeding the decarbonylative branch of the pathway and the other feeding the reductive branch. Thus, the *CER2* gene may affect the fatty acid elongation processes that feed the decarbonylative branch, and when this is blocked by the *cer2* mutation, fatty acid intermediates are diverted to the fatty acid elongation processes that feed the reductive branch, leading to increased primary alcohols.

### High Spatial Resolution Dynamics of Cuticular Surface Lipids

The larger accumulation of cuticular lipids on the abaxial side of the flower, primarily localized to the sepals, is consistent with the function of the cuticle as a water barrier (Aarts et al., 1995; Jung et al., 2006; Millar et al., 1999). Namely, because the sepals enclose the flower bud, their abaxial surfaces are outward facing, and thus the larger cuticular lipid load would provide more extensive protection from the desiccating environment.

Flowers transgenically expressing the *Gl2* or *Gl2-like* transgenes proved useful in visualizing the spatial distribution of the cuticular lipid biosynthetic reactions. This was accomplished by the co-localizing spectral images of substrate-product pairs of specific reactions of different segments of the cuticular lipid pathway. Thus, MSI analysis of flowers transgenically expressing the maize *Gl2* or *Gl2-like* genes demonstrate that the core processes of fatty acid elongation occur predominantly in the petals of flowers. Subsequently, the products of the FAE system are processed through the decarbonylative or reductive branch of the pathway. The co-localization spectral images of substrate-product pairs indicate that the decarbonylative branch is primarily expressed in the stigmatic tissue and sepals, and the reductive branch of the pathway is primarily expressed in the stigmatic tissue.

Therefore, the data presented in this study revealed previously unknown attributes concerning the spatial and temporal changes in cuticular lipid accumulation patterns, as affected by different genotypes and by flower development. These were revealed by the application of novel analytic technologies that directly assessed the accumulation of the metabolic intermediates and products of this complex pathway. Previously these were inferred by assessing cell-specific expression of genes and proteins associated with this pathway (Greer et al., 2007; Kim et al., 2019; Xia et al., 1996). Examples of these latter technologies include *in situ* hybridization to detect mRNA products of these genes or immunolocalization of protein gene-products, or by studies with reporter transgenes (e.g., GFP, RFP, GUS etc.) (Chalfie et al., 1994; Duck, 1994; Jefferson et al., 1987; Koo et al., 2007). Although these latter macromolecular analyses provide valuable insights on the gene expression program(s) that regulate cuticular lipid deposition, because of the complexity of the genetic and metabolic network that determines the accumulation of the final products, it’s difficult to dissect how these networks are regulated if one does not also have insights on the spatial and temporal patterns of the metabolic intermediate and products of the pathway. Hence, because complex pathways such as cuticular lipid biosynthesis, can integrate different regulatory processes that can be controlled by transcriptional, post-transcriptional or post-translational regulatory mechanisms, one needs a measure of the overall dynamics of cuticular lipid deposition, attributes that we have quantified in this study. Moreover, because this pathway is expressed by a single cell layer, it is also important to evaluate the heterogeneous spatial distribution of the products of the pathway across plant surfaces. In this study this heterogeneity was revealed relative to the different flower organs (i.e., sepals, petal, stigma and styles) and in relation to the abaxial and adaxial surfaces of these organs. Therefore, the metabolite imaging technique, coupled with specialized gas chromatography inlet that was used herein has proven to be a powerful combination for detecting the differential distribution and developmental changes in cuticular lipid deposition, providing new insights on the temporo-spatial granularity of the pathway.

In summary therefore, the application of these new analytical technologies that generated dynamic and high-spatially resolved data of the distribution of cuticular lipid metabolites has provided new insights into the cuticular lipid biosynthesis pathway. These include: i) a quantitative measure of the minimum flux through the pathway; ii) the finding that the reductive and decarbonylative branches of the pathway are spatially sequestered into separate flower organs; iii) the functional homology between the Arabidopsis *CER2* gene and the maize *Gl2* and *Gl2-like* genes; and iv) that the *cer2* mutation differentially affects the reductive and decarbonylative branches of the pathway on the pistil (i.e., stigma and style) and the perianth (i.e., petal and sepal) organs of the flower, respectively. Hence, these new insights reveal complexities in the spatial accumulation-patterns of the products of this pathway that will need more detailed biochemical and molecular dissection in order to mechanistically understand how the cuticular lipid biosynthesis pathway is spatially and temporally regulated.

## MATERIAL AND METHODS

### Plant Material and Growth Conditions

Arabidopsis genetic seed stocks were obtained from the Arabidopsis Biological Resource Center (www.arabidopsis.org). These included seeds of the wild-type Col-0 ecotype, and the SALK_084443C mutant line, which carried a T-DNA insertion mutation at the At4g24510 locus in the Col-0 background, i.e., the *cer2-5* mutant allele. Both these Arabidopsis lines used as hosts for transgenes that over-expressed the maize *Glossy2* (GRMZM2G098239; Zm00001d002353) or *Gl2-like* (GRMZM2G315767; Zm00001d024317) ORFs. In addition, we generated a control transgenic line that overexpressed the *CER2* ORF in the *cer2-5* mutant background. All these transgenes were constructed in the vector using pEarleyGate100 vector (Earley et al., 2006), which transcriptionally controlled the expression of the transgene with the CaMV 35S promoter. All plants were grown in a growth room maintained at 22 °C, and under continuous illumination, as described previously (Alexander et al., 2020). Genotypes of all transgenic plants were confirmed by PCR-based genotyping.

Biochemical analyses were conducted on Arabidopsis developing flowers (Figure 1) harvested at four defined stages of development: a) Stage A are flowers with closed buds; b) Stage B flowers were identified by the first emergence of petals from the bud; c) Stage C flowers were identified as presenting emerging petals that are perpendicular to the flower axis; d) Stage D flower reclosed and the developing silique was just starting to emerge. Time-lapse video-photography was used to determine the timing of the transitions between each stage of flower development, and weight of flowers was determined at each of the four development, from 2-3 biological replicates consisting of 6-13 pooled flowers.

### Cuticular lipid analysis of single Arabidopsis flowers

Extracellular cuticular lipids were extracted from individual flowers at each developmental stage (Stages A to D). Biologically replicated data were gathered by extracting extracellular cuticular lipids from six individual flowers at each stage of development, with each flower being harvested from independent plants. For quantification purposes, an aliquot of 0.01 μg hexacosane, dissolved in chloroform, was applied to the individual flowers, and after the solvent had evaporated, the flowers were immersed in 0.5-mL chloroform for 60s. After the flower had been removed, the chloroform solvent containing the extracted extracellular cuticular lipids, was dried under a stream of N_2_ gas. The dried extract was treated with 0.07-mL *N, O*-Bis (trimethylsilyl) trifluoroacetamide (BSTFA) with 1% trimethylchlorosilane (TMCS) (Sigma-Aldrich) 70°C for 30min, and 10μl was injected into the GC.

GC-MS analysis was performed with an Agilent 6890 GC interfaced to a 5973 mass spectrometer. The GC was equipped with a LVI-PTV inlet injector (Agilent Technologies, Santa Clara, CA), which enabled the analysis of dilute samples by providing a means of concentrating large injection volumes (10-μL). The GC was equipped with a HP-5ms column (30 m x 0.25 mm i.d. coated with a 0.25 μm film, Agilent Technologies). Chromatography was conducted with a mobile gas-phase of helium at a flow rate of 0.1 mL/min. The column oven temperature was programmed to increase from 80 °C to 180 °C at 20 °C/min, and held at this temperature for 1min, then ramped to 220 °C at a rate of 5°C/min, held at this temperature for 5min, and finally ramped to 320°C at 10°C/min, and held there for 10min. The interface temperature to the mass-spectrometer was at 280°C, and the ionization voltage for mass spectrometry was set at 70 eV.

The GC/MS data files were deconvoluted by National Institute of Standard and Technology Automated Mass spectral Deconvoluted and Identification System (NIST AMDIS) software, and searched against an in-house compound library as well as the NIST 14 Mass Spectral Library. The abundance of the extracellular lipids was calculated based on the peak size of the spiked hexacosane standard.

### Mass Spectrometric Imaging of flowers

Flowers collected at the four developmental stages were attached to Indium Tin Oxide (ITO) coated glass slides (Fisher Scientific, Pittsburg, PA) with double-sided tape. Flowers were carefully opened for imaging the adaxial surfaces of the floral organs using fine tweezers and a dissecting needle. For the abaxial (lower) surface, petals and sepals were carefully dissected and placed on the double-sided tape. Care was taken to ensure that all flower organs were flat on the MS imaging slide. The attached samples are dried in a glass vacuum desiccator for 30-60 min. Flowers were then sputter-coated with silver target 3NS (99.95% purity; ESPI Metals, Ashland, OR) for 50s, using a Cressington 308R Sputter Coater (Redding, CA, USA) at an argon partial pressure of 0.02 mbar, and a current of 80 mA.

A 7T SolariX Fourier Transform Mass Spectrometer (FTICR-MS) (BrukerDaltonics, Bremen, Germany) equipped with an MALDI ion source was used for MSI. The spectrometer used a SmartBeam™ II laser with a spot size of 80 μm diameter. All spectra were acquired in positive ion mode, integrating 200 laser shots per image pixel. The laser power was set to 25% with a frequency of 1000 Hz. The mass scanning range was set from *m/z* 300 to *m/z* 800, time of flight value was 0.7 s, and ion cooling time was set to 0.01 s. The mass spectrometer was externally calibrated prior to sample analysis, using arginine standard mixture (Sigma-Aldrich). Images were processed using FlexImaging 4.0 software (BrukerDaltonics, Bremen, Germany). Chemical identification of ions was performed based on theoretical monoisotopic masses of each surface lipid species and the mass tolerance was set to 2 ppm.

## ACKNOWLEDGEMENTS

The authors acknowledge Dr. Ann Perera, and Dr. Lucas J. Showman of the WM Keck Metabolomics Research Laboratory (Iowa State University, Ames, IA) for assistance in metabolomics analysis and mass spectral imaging; and Dr. Yue Wu (Iowa State University, Ames, IA) for the use of sputter coater equipment. This work was supported by the State of Iowa, through Iowa State University’s Center for Metabolic Biology, and by the National Science Foundation (grants IOS–1139489 and EEC–0813570 to B.J.N.), and the National Science Foundation-EAPSI program (grant OISE–1614020 to L.E.A.).

## AUTHOR CONTRIBUTIONS

B.J.N. conceived the project; L.E.A. generated and characterized transgenic lines, performed the single flower gas-chromatography experiments and analyzed the data; B.X., L.E.A., J.G. performed mass spectral imaging and data analysis and Z.S provided technical support in many aspects of the research; J.G. was an undergraduate research assistant supported by the SULI program of the Department of Energy Ames Laboratory; all authors contributed to the writing of the article.

## SUPPLEMENTAL MATERIAL

**Supplemental Figure 1.**
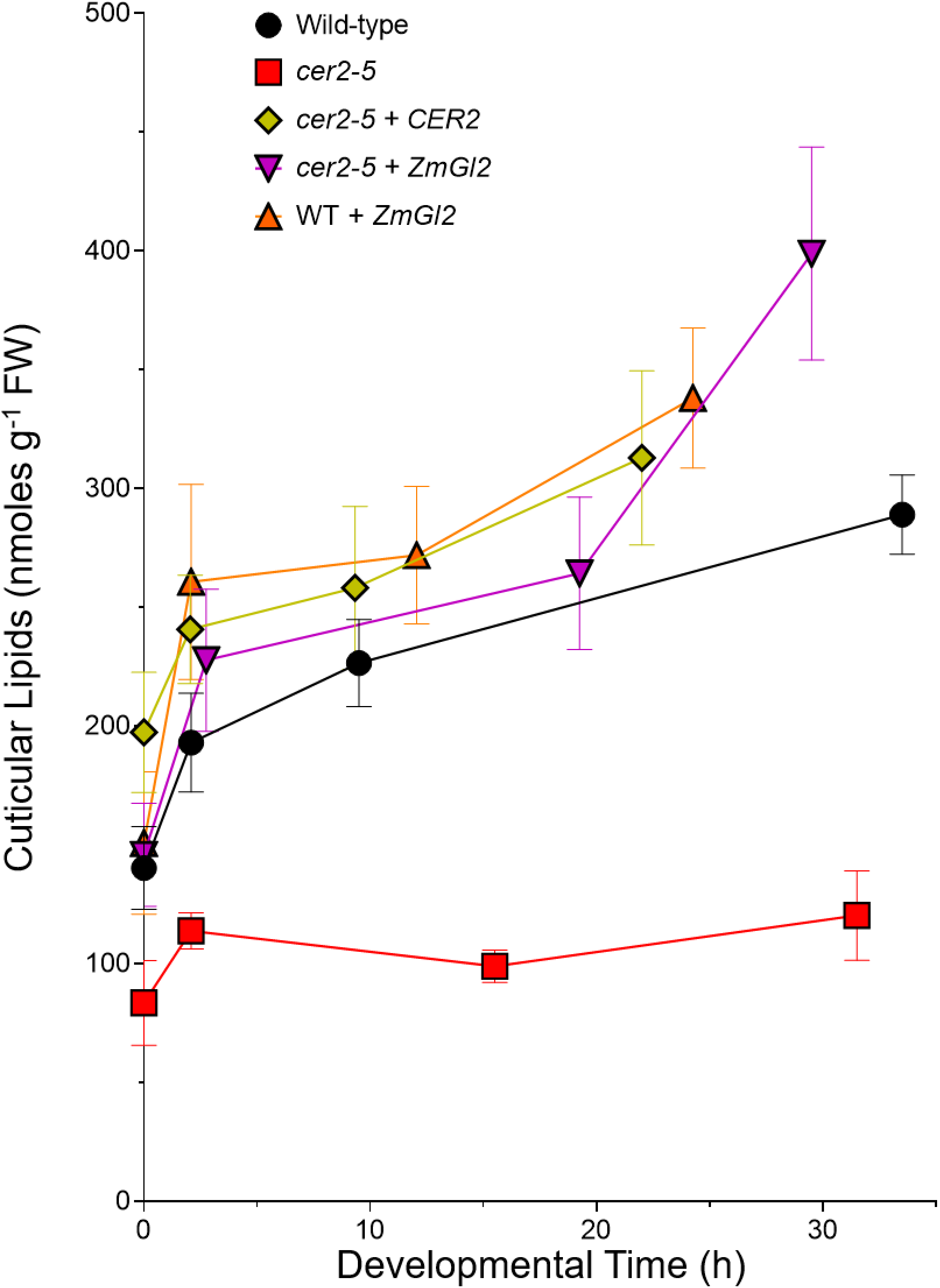
Effect of flower development on cuticular lipid accumulation. Cuticular lipids were extracted from individual Arabidopsis flowers of wild-type, *cer2-5* mutant, and either of these two backgrounds transgenically expressing the *CER2* ORF or the *ZmGl2* ORF. The extracts were analyzed by GC-MS, using a GC equipped with a LVI-PTV inlet injector (see Materials and Methods). The data represent average ± standard error of 6 replicates

**Supplemental Figure 2.**
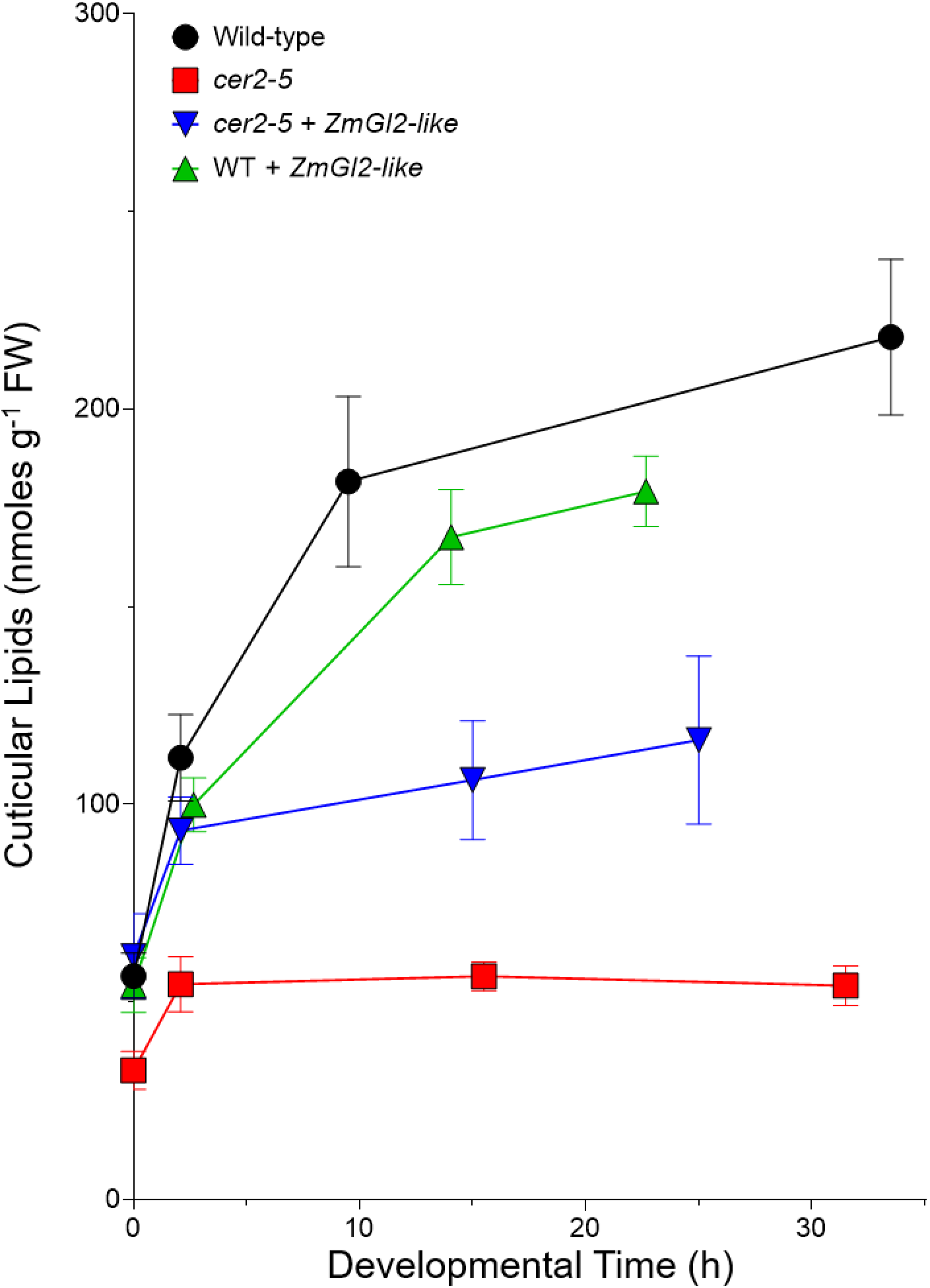
Effect of flower development on cuticular lipid accumulation. Cuticular lipids were extracted from individual Arabidopsis flowers of wild-type, *cer2-5* mutant, and either of these two backgrounds transgenically expressing the *ZmGl2-like* ORF. The extracts were analyzed by GC-MS, using GC equipped with a LVI-PTV inlet injector (see Materials and Methods). The data represent average + standard error of 6 replicates.

**Supplemental Figure 3.**
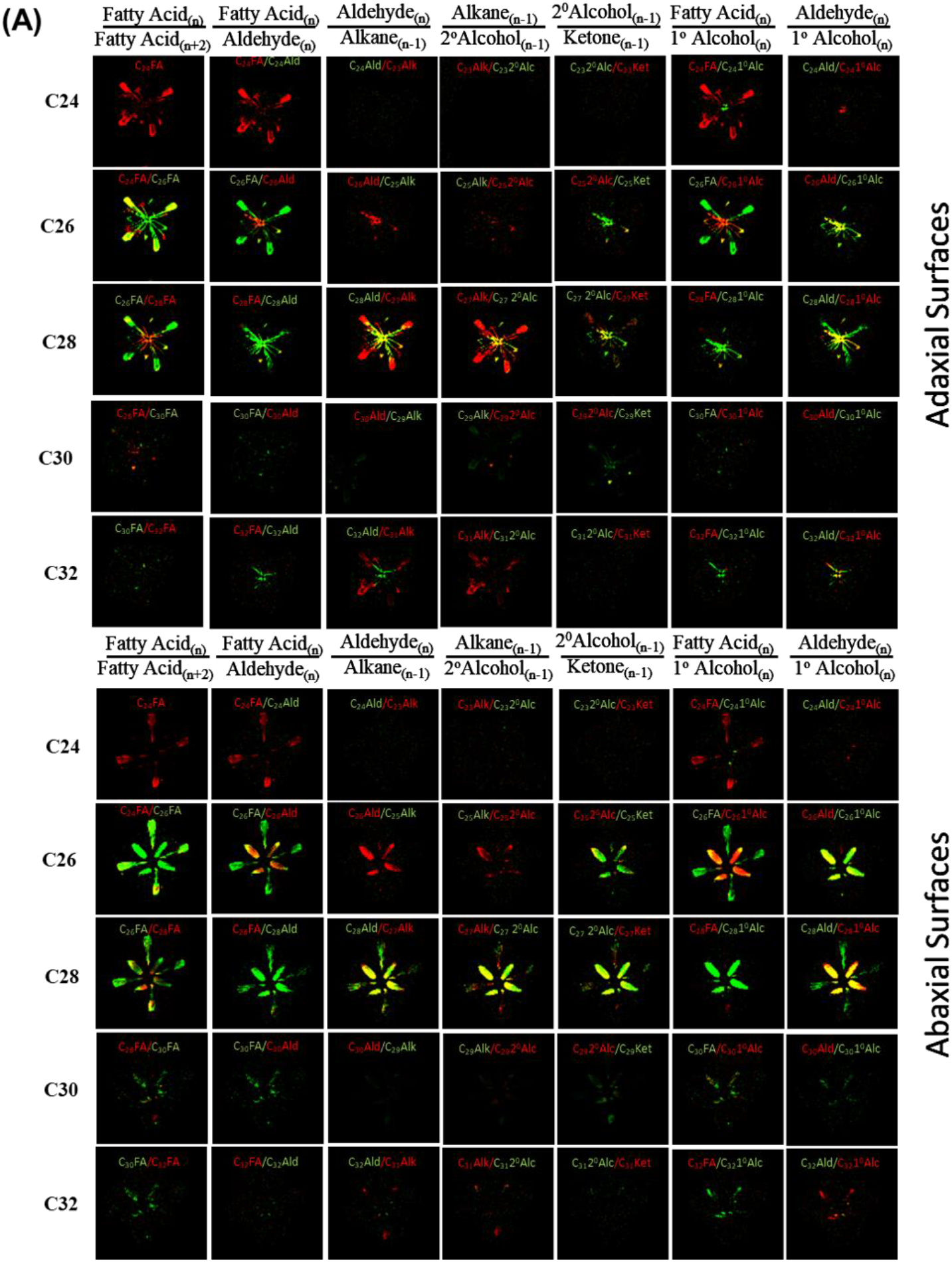
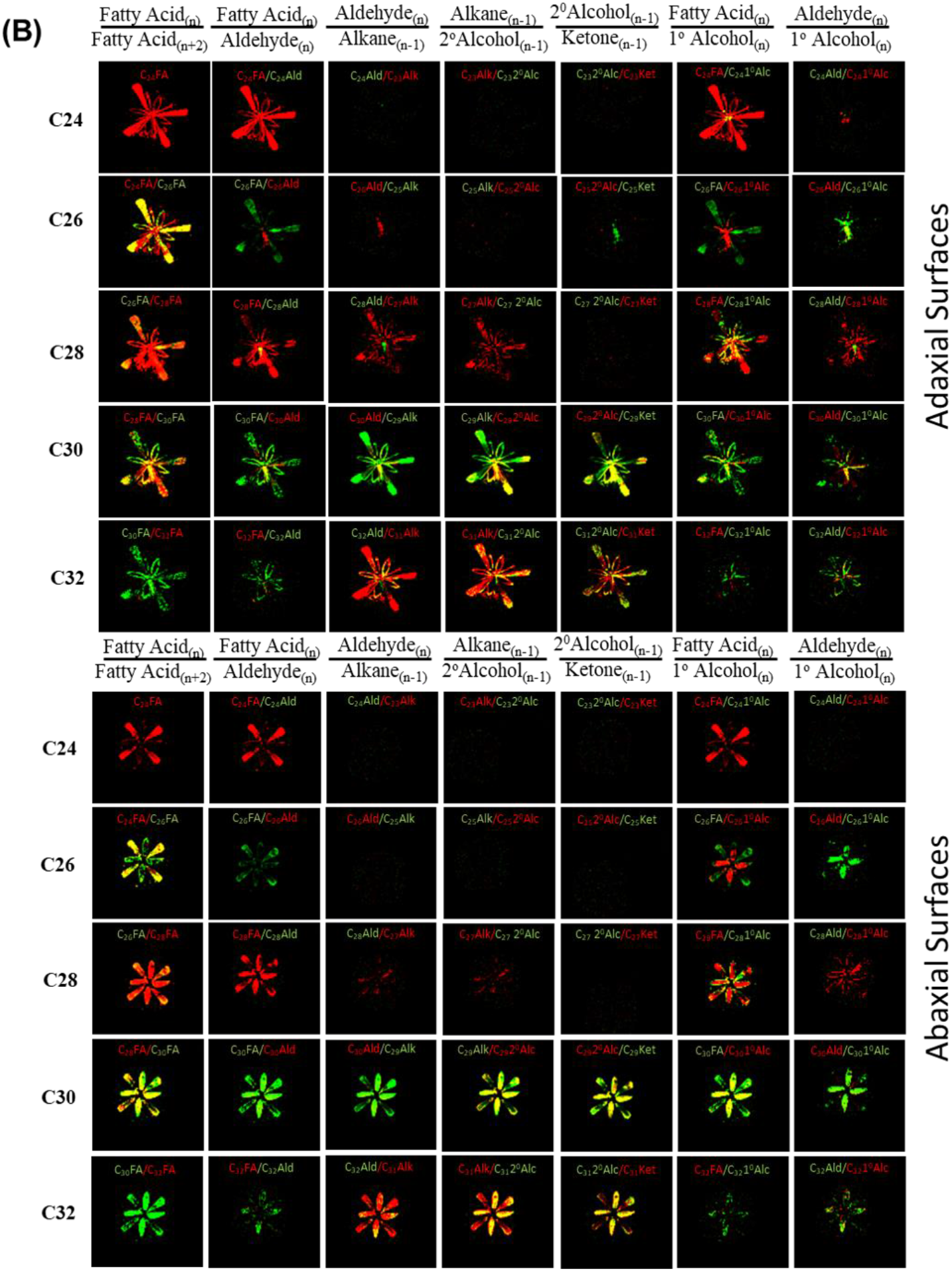
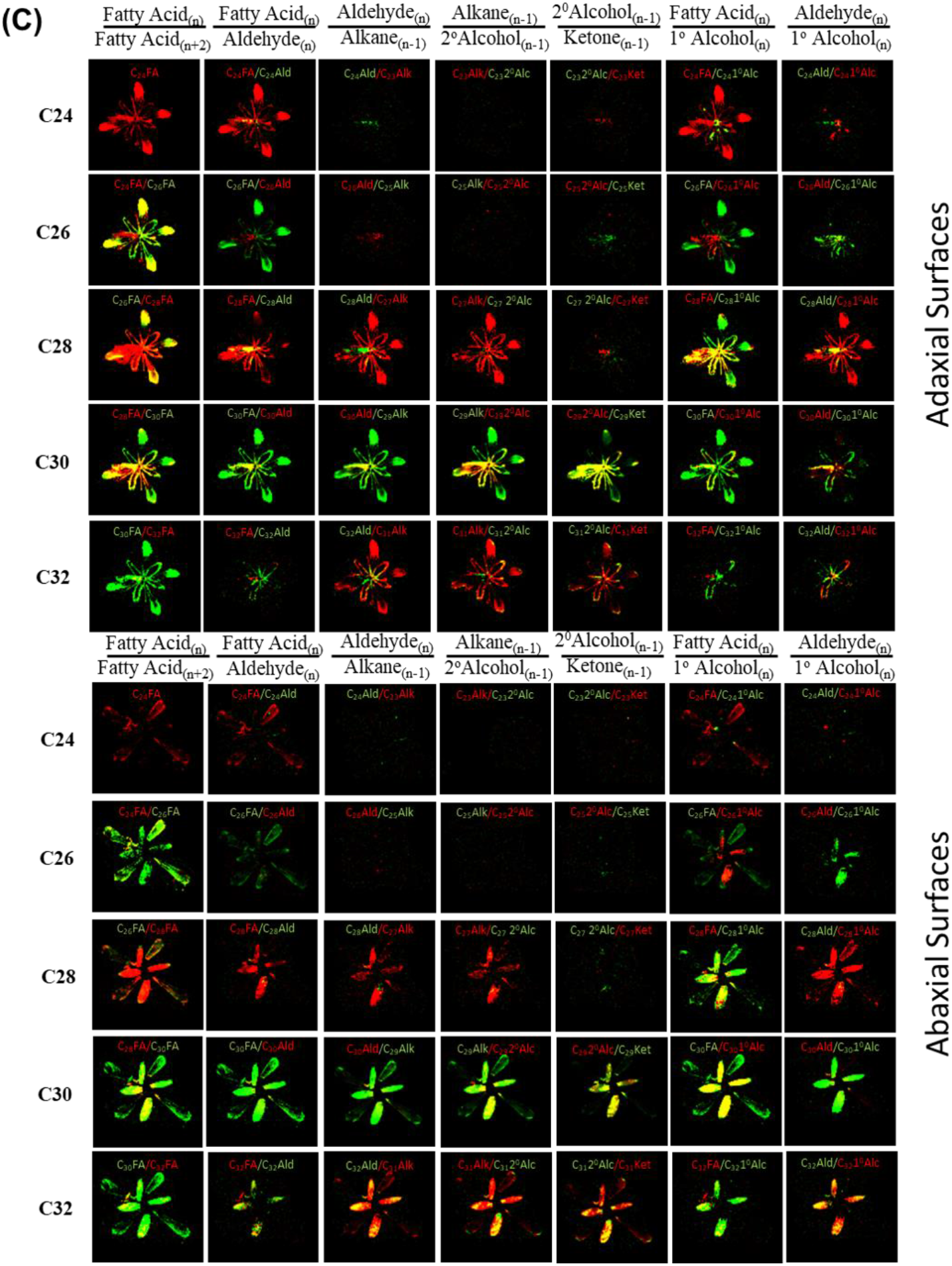
Mass spectrometric imaging of the co-localization of substrate-product pairs of individual reactions of the cuticular lipid biosynthesis pathway. Each image is a fusion of two false color-coded MSI images, green for the substrate, and red for the product of a single metabolic reaction that contributes to cuticular lipid biosynthesis pathway as denoted for 24:0 to 32:0 fatty acids and their alkyl derivatives. In these fused images, yellow represents the spatial zone where these substrate-product pairs co-localize, indicating the location of the underlying tissue/cells where this metabolic interconversion could be occurring. Images are indicated for the adaxial and abaxial surfaces of Arabidopsis flowers at developmental Stage C for the following genotypes: (A) *cer2-5* mutant, (B) *cer2-5* mutant transgenically expressing *Gl2-like*, (C) wild-type transgenically expressing *Gl2-like*.

**Supplemental Figure 4.**
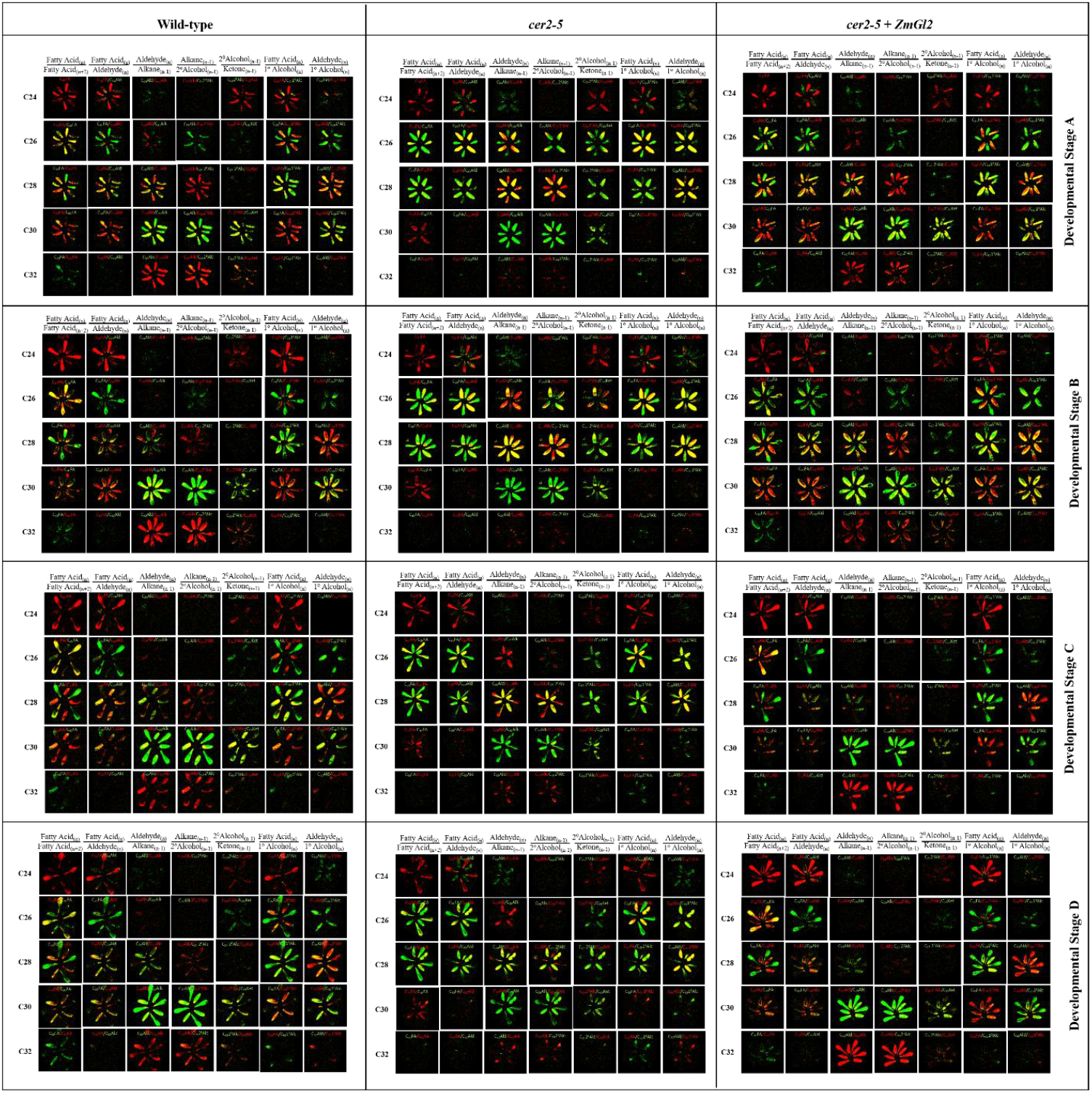
Mass spectrometric imaging of the co-localization of substrate-product pairs of individual reactions of the extracellular cuticular lipid biosynthesis pathway. Each image is a fusion of two false color-coded MSI images, green for the substrate, and red for the product of a single metabolic reaction that contributes to cuticular lipid biosynthesis pathway. In these fused images, yellow represents the spatial zone where these substrate-product pairs co-localize, indicating the location of the underlying tissue/cells where this metabolic interconversion could be occurring. Images are of the abaxial surfaces of Arabidopsis flowers, at developmental stages A-D of the indicated genotypes: wild-type; cer*2-5* mutant; and *cer2-5* mutant transgenically expressing *ZmGl2*

**Supplemental Figure 5.**
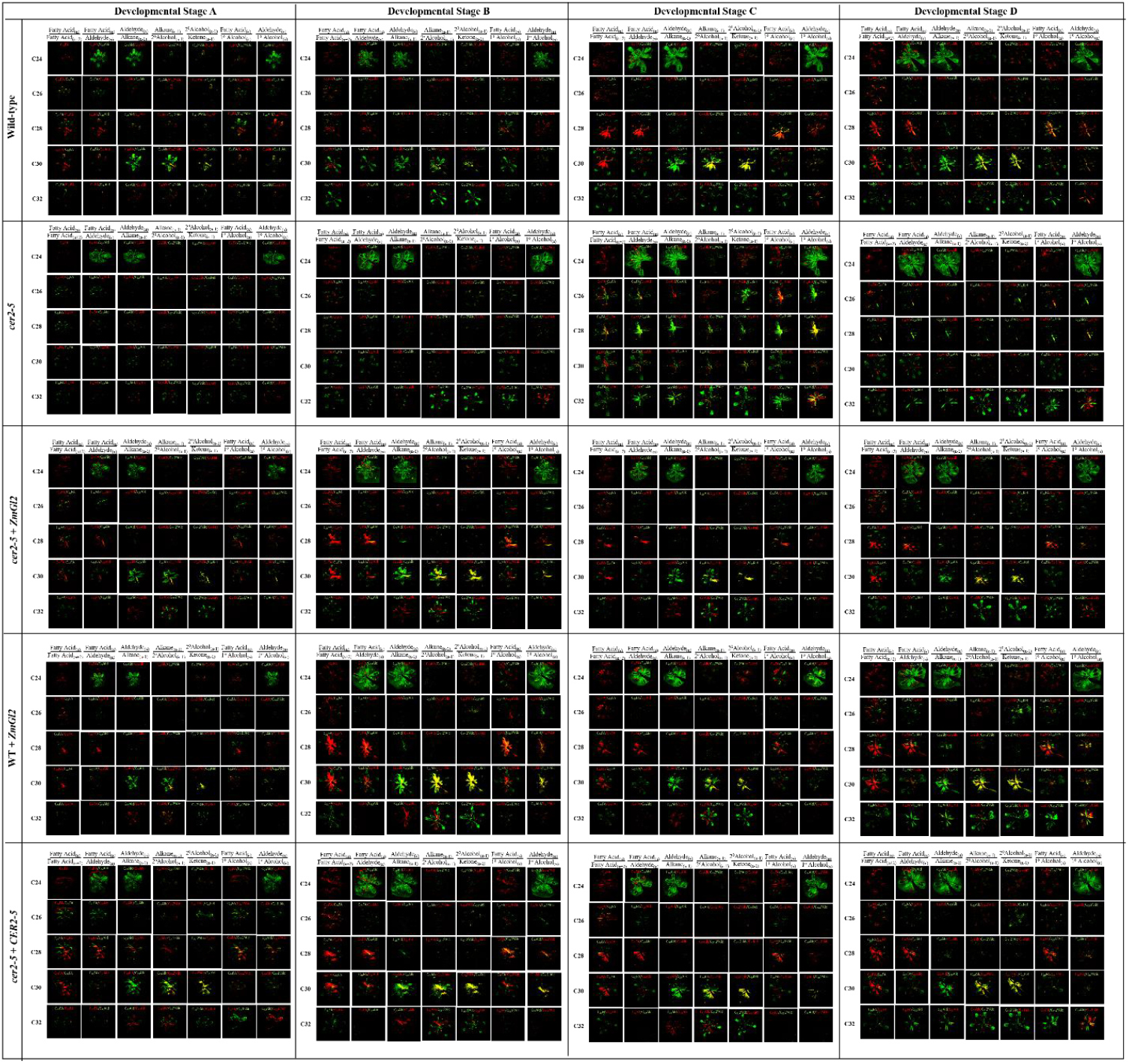
Mass spectrometric imaging of the co-localization of substrate-product pairs of individual reactions of the extracellular cuticular lipid biosynthesis pathway. Each image is a fusion of two false color-coded MSI images, green for the substrate, and red for the product of a single metabolic reaction that contributes to cuticular lipid biosynthesis pathway. In these fused images, yellow represents the spatial zone where these substrate-product pairs co-localize, indicating the location of the underlying tissue/cells where this metabolic interconversion could be occurring. Images are of the adaxial surfaces of Arabidopsis flowers, at developmental stages A-D of the indicated genotypes: wild-type; *cer2-5* mutant; *cer2-5* mutant transgenically expressing *ZmGl2*; wild-type transgenically expressing *ZmGl2*; and *cer2-5*mutant transgenically expressing *CER2*

**Supplemental Table 1.** Timing of flower development. Data were gathered from the indicated genotypes. The time to transition from developmental stage A to developmental stages B, C and D; average ± standard error (n = 2-4 replicates).

|  | Genotype |  |  |  |  |  |  |
| --- | --- | --- | --- | --- | --- | --- | --- |
|  | WT (Col-0) | *cer2-5* | *cer2-5* + *CER2* | *cer2-5* + *ZmGl2* | WT+*ZmGl2* | *cer2-5* + *ZmGl2*-like | WT+*ZmGl2*-like |
| Developmental Stage | Developmental Time (h) |  |  |  |  |  |  |
| Stage A | 00.00 ± 0.00 | 00.00 ± 0.00 | 00.00 ± 0.00 | 00.00 ± 0.00 | 00.00 ± 0.00 | 00.00 ± 0.00 | 00.00 ± 0.00 |
| Stage B | 02.09 ± 0.05 | 02.09 ± 0.09 | 02.09 ± 0.05 | 02.75 ± 0.48 | 02.09 ± 0.05 | 02.09 ± 0.05 | 02.67 ± 0.67 |
| Stage C | 09.50 ± 0.96 | 15.50 ± 0.50* | 09.33 ± 0.67* | 19.25 ± 3.09 | 12.04 ± 0.04* | 15.00 ± 4.04 | 14.06 ± 0.06* |
| Stage D | 33.50 ± 2.63 | 31.50 ± 4.50 | 22.00 ± 3.06 | 29.50 ± 3.77 | 24.25 ± 2.25* | 25.00 ± 6.35 | 22.67 ± 0.67* |
| Black and red asterisks denote time points that are significantly different from wild-type and *cer2-5* mutant, respectively, as determined by the statistical Student's T-test (p-value <0.05) |  |  |  |  |  |  |  |

|  | Genotype comparison |  |  |  |  |  |
| --- | --- | --- | --- | --- | --- | --- |
|  | *cer2-5* vs WT (Col-0) | *cer2-5* vs *CER2* in *cer2-5* | *cer2-5* vs *cer2-5*+*ZmGl2* | *cer2-5* vs *cer2-5*+*ZmGl2*-like | WT vs WT+*ZmGl2* | WT vs WT+*ZmGl2*-like |
| Developmental Stage | p-values determined by Student's T-test |  |  |  |  |  |
| Stage B | 1 | 0.725 | 0.408 | 1 | 1 | 0.347 |
| Stage C | 0.015 | 0.007 | 0.465 | 0.938 | 0.038 | 0.010 |
| Stage D | 0.699 | 0.164 | 0.767 | 0.548 | 0.037 | 0.019 |

**Supplemental Table 2.** Biomass of developing flowers. Data were gathered from the indicated genotypes. The data is the average of 6-13 pooled flowers at each stage of development, and estimated as the mean of 2-3 biological replicates ± standard deviation.

|  | Genotype |  |  |  |  |  |  |
| --- | --- | --- | --- | --- | --- | --- | --- |
|  | WT (Col-0) | *cer2-5* | *cer2-5+CER2* | *cer2-5+ZmGl2* | WT+*ZmGl2* | *cer2-5+ZmGl2-like* | WT+*ZmGl2-like* |
| Developmental Stage | Weight of Arabidopsis flowers (mg/flower) |  |  |  |  |  |  |
| Stage A | 0.63 ± 0.037 | 0.56 ± 0.095 | 0.40 ± 0.089 | 0.41 ± 0.078 | 0.44 ± 0.042* | 0.65 ± 0.050 | 0.69 ± 0.073 |
| Stage B | 0.73 ± 0.085 | 0.65 ± 0.059 | 0.47 ± 0.094 | 0.42 ± 0.028* | 0.50 ± 0.056* | 0.68 ± 0.021 | 0.71 ± 0.065 |
| Stage C | 0.84 ± 0.065 | 0.73 ± 0.128 | 0.59 ± 0.014 | 0.51 ± 0.016 | 0.72 ± 0.028 | 0.77 ± 0.069 | 0.82 ± 0.014 |
| Stage D | 0.94 ± 0.148 | 0.81 ± 0.097 | 0.62 ± 0.003 | 0.50 ± 0.049* | 0.72 ± 0.021 | 0.95 ± 0.012 | 1.04 ± 0.186 |
| Black and red asterisks denote flowers whose biomass are significantly different from wild-type and *cer2-5* mutant, respectively, as determined by the statistical Student's T-test (p-value <0.05) |  |  |  |  |  |  |  |

|  | Genotype comparison |  |  |  |  |  |
| --- | --- | --- | --- | --- | --- | --- |
|  | *cer2-5* vs WT (Col-0) | *cer2-5* vs *CER2* in *cer2-5* | *cer2-5* vs *cer2-5+ZmGl2* | *cer2-5* vs *cer2-5+ZmGl2-like* | WT vs WT+*ZmGl2* | WT vs WT+*ZmGl2-like* |
| Developmental Stage | p-value as determined by Student's T-test |  |  |  |  |  |
| Stage A | 0.314 | 0.147 | 0.150 | 0.316 | 0.013 | 0.303 |
| Stage B | 0.271 | 0.068 | 0.015 | 0.461 | 0.047 | 0.519 |
| Stage C | 0.264 | 0.233 | 0.104 | 0.710 | 0.098 | 0.545 |
| Stage D | 0.288 | 0.073 | 0.026 | 0.160 | 0.136 | 0.558 |

**Supplemental Table 3.** Rates of cuticular lipid accumulation during development of Arabidopsis flowers. Data were gathered from the indicated genotypes, at the initial stage of flower development (transition from developmental stages A to B), and the later stage of flower development (transition from developmental stages C and D). The data is the average gathered from 6 individual flowers at each stage of development.

| Metabolite Class | Alkyl chain length | Genotypes |  |  |  |  |  |  |  |
| --- | --- | --- | --- | --- | --- | --- | --- | --- | --- |
|  |  | Wild-type (Col-0) |  | *cer2-5* |  | *cer2-5* + *ZmGl2-like* |  | WT + *ZmGl2-like* |  |
|  |  | Stages A-B transition | Stages C-D transition | Stages A-B transition | Stages C-D transition | Stages A-B transition | Stages C-D transition | Stages A-B transition | Stages C-D transition |
|  |  | Rate of accumulation (nmol h<sup>-1</sup> g<sup>-1</sup> FW) |  |  |  |  |  |  |  |
| Fatty Acids | 20 | 0.019 | 0.003 | 0.024 | 0.010 | 0.015 | 0.007 | -0.011 | 0.008 |
|  | 22 | 0.019 | 0.001 | 0.010 | 0.004 | 0.015 | 0.002 | 0.006 | 0.006 |
|  | 24 | 0.106 | -1E-04 | 0.130 | -0.003 | 0.148 | -0.014 | 0.024 | -0.017 |
|  | 26 | 0.044 | 0.002 | 0.133 | 0.015 | 0.040 | -0.002 | 0.003 | 0.003 |
|  | 28 | -3E-05 | -1E-06 | 0.174 | 0.015 | 0.040 | -0.010 | -4E-06 | -9E-06 |
|  | 30 | -3E-05 | 0.001 | -3E-05 | -2E-06 | -8E-06 | 3E-04 | -4E-06 | 0.003 |
| Primary Alcohol | 24 | 0.012 | 0.001 | 0.006 | 0.002 | 0.030 | -0.003 | 0.011 | 0.004 |
|  | 26 | 0.109 | 0.014 | 0.478 | 0.116 | 0.388 | 0.071 | 0.057 | 0.021 |
|  | 28 | 0.264 | 0.037 | 0.409 | 0.178 | 0.319 | 0.065 | 0.166 | 0.041 |
|  | 30 | 0.112 | 0.017 | 1E-04 | 7E-05 | 0.054 | 0.012 | 0.111 | 0.007 |
| Aldehyde | 30 | -3E-05 | 0.008 | -3E-05 | -2E-06 | 0.034 | 0.003 | -0.021 | 0.011 |
| Alkane | 25 | -0.021 | -0.003 | -0.046 | -0.014 | -0.017 | -0.024 | -0.006 | -0.025 |
|  | 27 | 0.976 | -0.026 | 2.148 | -0.113 | 0.696 | -0.117 | 0.348 | -0.112 |
|  | 29 | 17.513 | 0.923 | 1.978 | -0.247 | 8.784 | 0.700 | 12.067 | 0.923 |
|  | 31 | 0.422 | 0.011 | 0.214 | -0.014 | 0.375 | 0.023 | 0.845 | 0.013 |
| Sec. Alcohol | 27 | -3E-05 | -1E-06 | 0.025 | 0.007 | 0.009 | 0.003 | -4E-06 | -9E-06 |
|  | 29 | 1.730 | 0.230 | 0.003 | 2E-04 | 0.518 | 0.228 | 1.219 | 0.390 |
| Ketone | 29 | 1.223 | 0.313 | 0.014 | 0.002 | 0.455 | 0.340 | 0.875 | 0.481 |

**Supplemental Table 4.** Rates of cuticular lipid accumulation during development of Arabidopsis flowers. Data were gathered from the indicated genotypes at the initial stage of flower development (transition from developmental stages A and B) and the latter stage of flower development (transition from developmental stages C and D). The data is the average gathered from 6 individual flowers at each stage of development.

| Metabolite Class | Alkyl chain length | Genotypes |  |  |  |  |  |  |  |  |  |
| --- | --- | --- | --- | --- | --- | --- | --- | --- | --- | --- | --- |
|  |  | Wild-type (Col-0) |  | *cer2-5* |  | *cer2-5* + *ZmGl2* |  | WT + *ZmGl2* |  | *cer2-5* + *CER2* |  |
|  |  | Stages A-B transition | Stages C-D transition | Stages A-B transition | Stages C-D transition | Stages A-B transition | Stages C-D transition | Stages A-B transition | Stages C-D transition | Stages A-B transition | Stages C-D transition |
|  |  | Rate of accumulation (nmol h<sup>-1</sup> g<sup>-1</sup> FW) |  |  |  |  |  |  |  |  |  |
| Fatty Acids | 20 | 0.161 | 0.004 | -0.059 | -0.002 | 0.005 | 0.019 | -0.003 | 0.024 | 0.618 | -0.015 |
|  | 24 | 0.590 | 0.014 | 0.405 | -0.054 | 0.581 | 0.063 | 0.725 | 0.092 | 1.155 | -0.055 |
|  | 26 | 0.185 | 0.003 | 0.231 | -0.029 | 0.162 | 0.018 | 0.039 | 0.018 | 0.743 | -0.082 |
| Primary Alcohol | 22 | -0.021 | -0.029 | 0.013 | 0.009 | 0.190 | 0.273 | -0.005 | 0.097 | 0.386 | -0.002 |
|  | 26 | 0.003 | 0.037 | 0.597 | 0.594 | 0.071 | 0.432 | 0.144 | 0.047 | 0.195 | 0.212 |
|  | 28 | 0.106 | 0.089 | 1.935 | 0.663 | 0.173 | 0.459 | 0.247 | 0.091 | 0.546 | 0.689 |
|  | 30 | 0.027 | 0.018 | -0.004 | 0.004 | 0.107 | 0.236 | 0.152 | 0.044 | 0.271 | 0.075 |
| Alkane | 25 | 0.306 | -0.012 | -0.246 | -0.041 | -0.106 | -0.078 | 0.063 | -0.013 | 0.174 | 0.035 |
|  | 27 | 0.598 | -0.037 | 2.168 | -0.005 | 0.387 | -0.044 | 0.466 | -0.097 | 0.969 | 0.032 |
|  | 29 | 11.882 | 1.322 | 2.509 | 0.097 | 13.793 | 5.592 | 24.136 | 2.186 | 4.442 | 1.944 |
|  | 31 | 1.684 | 0.155 | -0.014 | -0.035 | 3.964 | 1.838 | 12.689 | 0.926 | 1.163 | 0.019 |
| Sec. Alcohol | 29 | 0.982 | 0.488 | 0.024 | 0.008 | 1.377 | 1.597 | 2.729 | 0.577 | 0.753 | 0.551 |
| Ketone | 29 | 1.319 | 0.505 | 0.011 | 0.010 | 1.289 | 1.954 | 3.213 | 0.957 | 1.753 | 0.638 |

**Supplemental Data Sheet 1.** Raw data of cuticular lipid analysis extracted from single flowers.

